# Proton-coupled alternating access in a versatile mycobacterial Spns drug transporter

**DOI:** 10.64898/2026.05.09.724020

**Authors:** Kevin L. Jagessar, Samantha Gies, Tianqi Wu, Ian Miller, Khadijeh Dastvan, Reza Dastvan

**Affiliations:** Department of Biochemistry and Molecular Biology, Saint Louis University School of Medicine, St. Louis, MO, USA; Center for Applied AI for Protein Dynamics, Vanderbilt University, Nashville, TN, USA

**Author notes:** These authors contributed equally to this work (Kevin L. Jagessar, Samantha Gies). These authors contributed equally to this work (Tianqi Wu, Ian Miller).

## Abstract

Spns transporters are a mechanistically distinct branch of the major facilitator superfamily that regulate lipid transport, lysosomal homeostasis, immunity and disease. How the conserved Spns fold integrates protonation, substrate binding and alternating access to support chemically and directionally diverse transport remains unresolved. Here we combine DEER spectroscopy in lipid nanodiscs with DEER-and AlphaFold-guided modeling and protonation-mimetic mutagenesis to define the conformational landscape of the *Mycobacterium smegmatis* homolog *Ms*Spns. Protonation shifts *Ms*Spns toward an inward-facing state, whereas deprotonation favors a broader outward-facing ensemble through remodeling of intracellular and extracellular gates. Protonation-mimetic substitutions identify Glu126 as a switch that stabilizes an inward-facing, substrate-entry-competent conformation, while Asp38 and Asp57 favor outward-facing states and tune the extracellular proton-sensing network. The substrate-binding cavity displays distinct proton sensitivity and weaker cooperativity than gating networks. Hydrophilic cationic substrates stabilize the outward-facing state, consistent with efflux antiport, whereas lipophilic compounds favor the inward-facing state, suggesting uptake or allosteric stabilization. Thus, conserved proton-coupling elements can power opposing transport modes, revealing the mechanistic versatility of the Spns fold and its therapeutic potential.

---

Spns transporters have recently emerged as a mechanistically distinct and physiologically important branch of the major facilitator superfamily (MFS), with central roles in lipid transport, lysosomal homeostasis, immunity, and human disease^1–10^. The human Spns family comprises three lipid transporters, Spns1-3. Spns2 exports the bioactive lysolipid sphingosine-1-phosphate (S1P) from endothelial cells, thereby regulating S1P levels in circulatory fluids, lymphocyte trafficking, vascular function, and cell survival, and positioning Spns2 as a compelling immunomodulatory drug target^11^. Structural studies have rapidly transformed this field. Cryo-EM analyses captured human Spns2 in inward-facing (IF), occluded (O), and outward-facing (OF) conformations and revealed a noncanonical alternating-access cycle, while also defining the structural basis of inhibition^2^. Subsequent structures of Spns2 bound to S1P, inhibitors, and the immunomodulator FTY720-P further delineated the substrate-binding cavity, identified residues required for cargo recognition and conformational coupling, and provided an increasingly complete framework for transport and pharmacological modulation^3,4,6^. In addition, PI(4,5)P_2_ acts as a synergistic regulator that amplifies S1P-induced gate dynamics and transport^5^.

Progress on Spns1 has similarly broadened the biological scope of the family. Initially linked to lysosomal function and autophagic lysosome reformation, Spns1 is now recognized as a lysosomal lysophospholipid exporter that salvages phospholipids from the lysosomal lumen to the cytosol^8–10^. Functional studies established that Spns1 mediates the rate-limiting efflux of lysophosphatidylcholine (LPC) and lysophosphatidylethanolamine (LPE) and is essential for lysosomal lipid homeostasis^8^. This assignment is reinforced by mouse studies showing that Spns1 loss causes lysolipid accumulation and lysosomal storage disease-like phenotypes^9^. More recently, a cryo-EM structure of human Spns1 in an LPC-bound lumen-facing state identified structural features governing lysophospholipid recognition, revealed a TM5-TM8 luminal gate specialized for substrate entry, and uncovered a residue network implicated in proton sensing^10^. Consistent with this physiological importance, biallelic Spns1 loss-of-function variants cause a multiorgan disorder marked by liver and muscle injury, lysosomal lipid accumulation, and disrupted mTOR-regulated lipid homeostasis^12^. Together, these findings highlight the biomedical importance of defining how the Spns fold couples protonation, substrate chemistry, membrane lipids, and alternating access.

Despite these advances, key aspects of the transport mechanism remain unresolved, including energy coupling and how conserved Spns proton-coupling elements reshape the conformational landscape to enable chemically diverse and directionally distinct transport modes. Using an integrated approach in lipid membranes, we previously defined proton-and substrate-coupled conformational dynamics in the bacterial Spns homolog from *Hyphomonas neptunium* (*Hn*Spns)^7,13^. We identified conserved residues that regulate protonation and showed how sequential protonation of proton-switch residues coordinates conformational transitions. Consistent with this model, some recent studies support proton-coupled transport in Spns2^3^, whereas others propose alternative coupling mechanisms^2,14^ or emphasize substrate binding, conformational states, lipid regulation, or inhibition. Recent work on Spns1^8,10^ also strongly supports proton-coupled transport. Although the proposed luminal proton-sensing network in Spns1 is supported by mutational and cell-based transport studies, the contribution of membrane-embedded proton-sensing networks remains unexplored. Likewise, reverse mutation of the corresponding luminal network in Spns2 restores pH dependence of S1P transport^10^, but this does not exclude additional extracellular proton-sensing networks in Spns2 or other Spns proteins.

The bacterial Spns transporter MSMEG_3705 from *Mycobacterium smegmatis* (*Ms*Spns) provides a powerful system to address this gap by linking bacterial efflux and influx physiology to broader mechanistic principles governing lipid-transporting MFS proteins. *Ms*Spns has been implicated in selective drug resistance and broader mycobacterial physiology^7,15^. The structurally unrelated but hydrophilic antimicrobial compounds ethidium bromide and capreomycin, a cyclic polypeptide antituberculosis antibiotic, are substrates of *Ms*Spns, whereas the transporter does not confer resistance to the highly lipophilic first-line antituberculosis drug rifampicin^7,15^. Deletion of *Ms*Spns increases intracellular ethidium bromide accumulation, enhances capreomycin sensitivity, reduces rifampicin susceptibility, and accelerates growth in association with upregulation of the downstream isocitrate lyase gene^15^. These phenotypes suggest that *Ms*Spns couples selective efflux, possible uptake-like behavior, and adaptive mycobacterial physiology.

Spns proteins are secondary-active MFS transporters that use cation electrochemical gradients, most notably protons, to drive substrate transport^16–18^. Like other MFS transporters, they contain 12 transmembrane helices (TMs) arranged into two pseudosymmetrical six-helix domains, the N-terminal domain (NTD, TMs 1-6) and the C-terminal domain (CTD, TMs 7-12), which enclose a central substrate-binding and translocation cavity (Fig. 1)^7,19,20^. These helices form four three-helix repeats, with TMs 1, 4, 7, and 10 lining the cavity, TMs 2, 5, 8, and 11 shaping the lateral gates, and TMs 3, 6, 9, and 12 providing structural support at the membrane interface^16^ (Fig. 1c). The NTD is more strongly conserved across bacterial and eukaryotic Spns transporters, whereas the CTD is more divergent, suggesting a conserved mechanistic core superimposed on lineage-specific functional specialization (Figs. 1b, 1d–f)^7^. Sequence alignments and residue maps of *Ms*Spns and human Spns2 reveal highly conserved clusters of charged and polar residues within the transmembrane core, around the central cavity, and at the intracellular and extracellular gating interfaces (Fig. 1), consistent with an extensive charge-relay and proton-coupling architecture. These residues include putative membrane-embedded protonation switches (Fig. 1b), intradomain charge-relay interactions (Fig. 1d–f), and an extracellular proton-sensing network (Fig. 1a), all positioned to couple protonation to alternating access. Defining how protonation remodels this network is therefore central to understanding transport across the Spns family.

**Figure 1.**
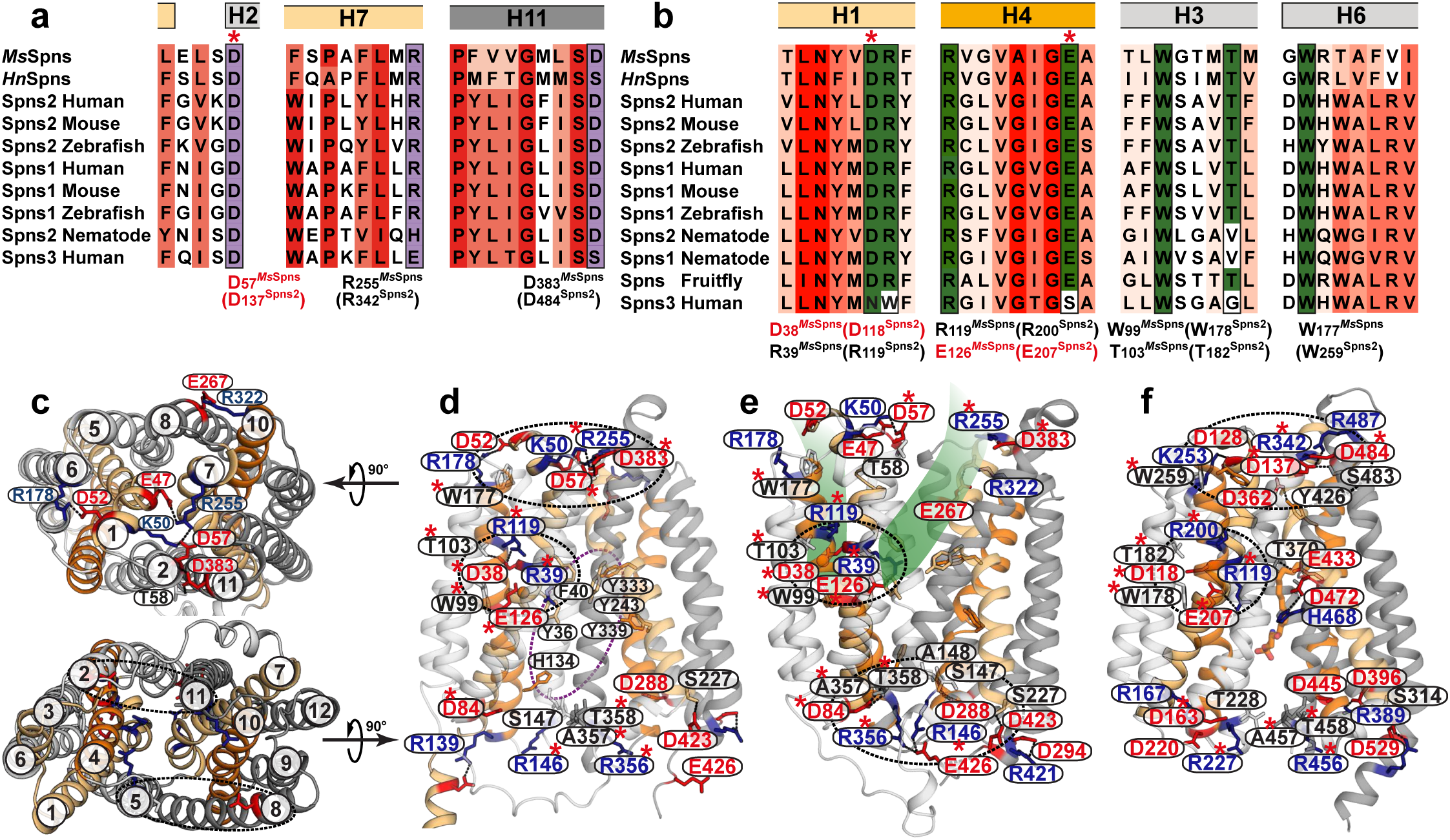
Domain architecture and conserved functional residues of Spns proteins. (**a**) Sequence alignment highlighting conserved elements of the extracellular proton-sensing network, shaded in purple. (**b**) Sequence alignment highlighting conserved residues of the membrane-embedded protonation switches and the putative periplasmic proton-translocation pathway, shaded in green. (**c**) Periplasmic view of the inward-facing (IF) *Ms*Spns model showing the proton-sensing network as sticks, and cytoplasmic view showing the N-terminal domain (NTD) and C-terminal domain (CTD) in light and dark grey, respectively. (**d,e**) Membrane views of IF (**d**) and outward-facing (OF; **e**) *Ms*Spns, highlighting conserved Spns-family residues marked with asterisks and *Ms*Spns-specific functional residues. Symmetry-related TM helices 1/7 and 4/10 are colored light and dark orange, respectively. Membrane-embedded protonation switches, extracellular and intracellular proton-sensing networks, and conserved aromatic cavity residues are indicated by circles. (**f**) Cryo-EM structure of human Spns2 bound to S1P (Protein Data Bank [PDB] code 8EX4), highlighting functionally conserved residues in mammalian Spns proteins.

Here, we combine comprehensive double electron-electron resonance (DEER; also called PELDOR) spectroscopy^7,21–23^ in lipid nanodiscs with DEER-and AlphaFold-guided molecular modeling^24–27^ to define the conformational landscape of *Ms*Spns and determine how protonation, protonation-mimetic mutations, ligand chemistry, and membrane lipids bias its energy landscape. By integrating pH-dependent distance measurements with protonation-mimetic substitutions, we directly test how conserved protonation switches and proton-sensing residues control intracellular gating, extracellular closure, and substrate-cavity remodeling, thereby revealing both shared and divergent mechanisms across the Spns family. Our results establish the molecular basis for the dual behavior of this mycobacterial Spns transporter, including export of cationic hydrophilic compounds and uptake-like stabilization by lipophilic ligands. More broadly, they show how the conserved Spns fold can support bacterial efflux and influx while preserving core proton-coupling principles that link *Ms*Spns to the emerging structural and dynamic framework of mammalian Spns transporters.

## Results

Mechanistic dissection of the Spns transport cycle requires defining the conformational states associated with substrate binding, translocation and release, and the protonation switches that drive transitions between them. Because the cellular proton gradient is inwardly directed, *Ms*Spns could operate either as a proton symporter that mediates substrate uptake or as an antiporter that couples proton influx to substrate efflux. To define its proton and ligand dependence, we measured DEER distance distributions at pH 4 to favor protonation, pH 7.5 to represent neutral conditions, and pH 9 to favor deprotonation. We also combined these pH-dependent measurements with protonation-mimetic substitutions of conserved acidic residues, D38N, E126Q, D38N-E126Q and D57N (Figs. 1a and 1b), to test whether protonation-mediated charge neutralization at these sites is sufficient to bias *Ms*Spns toward defined conformational states. For selected reporter pairs spanning the substrate-binding cavity and gating helices, we also measured conformational responses to confirmed and putative substrates^7,15^. All experiments were performed with *Ms*Spns reconstituted into *E. coli* polar lipid nanodiscs, with additional lipid-composition comparisons for selected pairs.

### Structural and functional integrity of *Ms*Spns mutants

For DEER measurements, cysteine substitutions were introduced at selected sites on a cysteine-less (CL) background. We assessed the functional integrity of the CL construct and unlabeled double-cysteine mutants relative to wild-type (WT) *Ms*Spns using a previously established *Escherichia coli* cell-growth assay that reports resistance to toxic capreomycin concentrations and serves as a surrogate for *Ms*Spns-mediated drug efflux^28^. Consistent with our earlier findings that *Ms*Spns confers resistance to capreomycin and ethidium bromide, but not rifampicin^7,15^, cells expressing the CL and double-cysteine mutants survived capreomycin concentrations that were lethal to cells carrying the empty vector (Supplementary Fig. 1), confirming that the engineered constructs retained function.

As an additional probe of structural integrity, we monitored low-pH fluorescence quenching in spin-labeled *Ms*Spns reconstituted into lipid nanodiscs as a surrogate readout of conformational change (Supplementary Fig. 2)^29^. Protonation of WT *Ms*Spns at pH 4 reduced Trp fluorescence relative to pH 9, consistent with a protonation-dependent change in the local environment of Trp residues. Statistical analysis showed no significant differences between the spin-labeled double-cysteine constructs and CL *Ms*Spns, indicating that cysteine introduction and spin labeling at the probed sites did not measurably perturb this conformational response. Together, these assays support the structural and functional integrity of the reporter constructs, although direct effects of spin labeling on transport activity could not be assessed.

### Protonation promotes opening of the intracellular side

Motif A, the most conserved sequence motif in MFS transporters, lies in the intracellular loop between TM2 and TM3 (Fig. 2b)^16,30^. Together with surrounding residues from TM4 and TM11, it forms the conserved structural motif A, which stabilizes the O or OF state^16^. In Spns2 and Spns1, an associated charge-relay network (Asp163^TM2^-Arg167^TM3^-Asp220^TM4^ in Spns2) is proposed to regulate the interdomain charge-dipole interaction between TMs 2 and 11 (Fig. 2b). In mammalian Spns proteins, a symmetry-related motif A-like network is also formed by TMs 8, 9, 10, and 5 (Asp396^TM9^-Arg389^TM8^-Asp445^TM10^-Arg227^TM5^ in Spns2). These canonical motif A interactions are only partially conserved in *Ms*Spns. Backbone and side-chain hydrogen-bonding interactions between TM2 and TM11 (Asp84^TM2^-Ala357^TM11^/Thr358^TM11^) and between TM8 and TM5 (Asp288^TM8^-Ser147^TM5^/Ala148^TM5^) remain intact (Fig. 2a). However, *Ms*Spns uniquely lacks the highly conserved motif A aspartates on TMs 4 and 10 and therefore cannot form the corresponding interdomain salt bridges with Arg356^TM11^ and Arg146^TM5^, respectively (Fig. 2b). In the AlphaFold3-derived OF model of *Ms*Spns, these missing interactions appear to be compensated by an alternative Arg356^TM11^:Glu426^TM12^:Arg146^TM5^ salt-bridge network (Fig. 2a).

**Figure 2.**
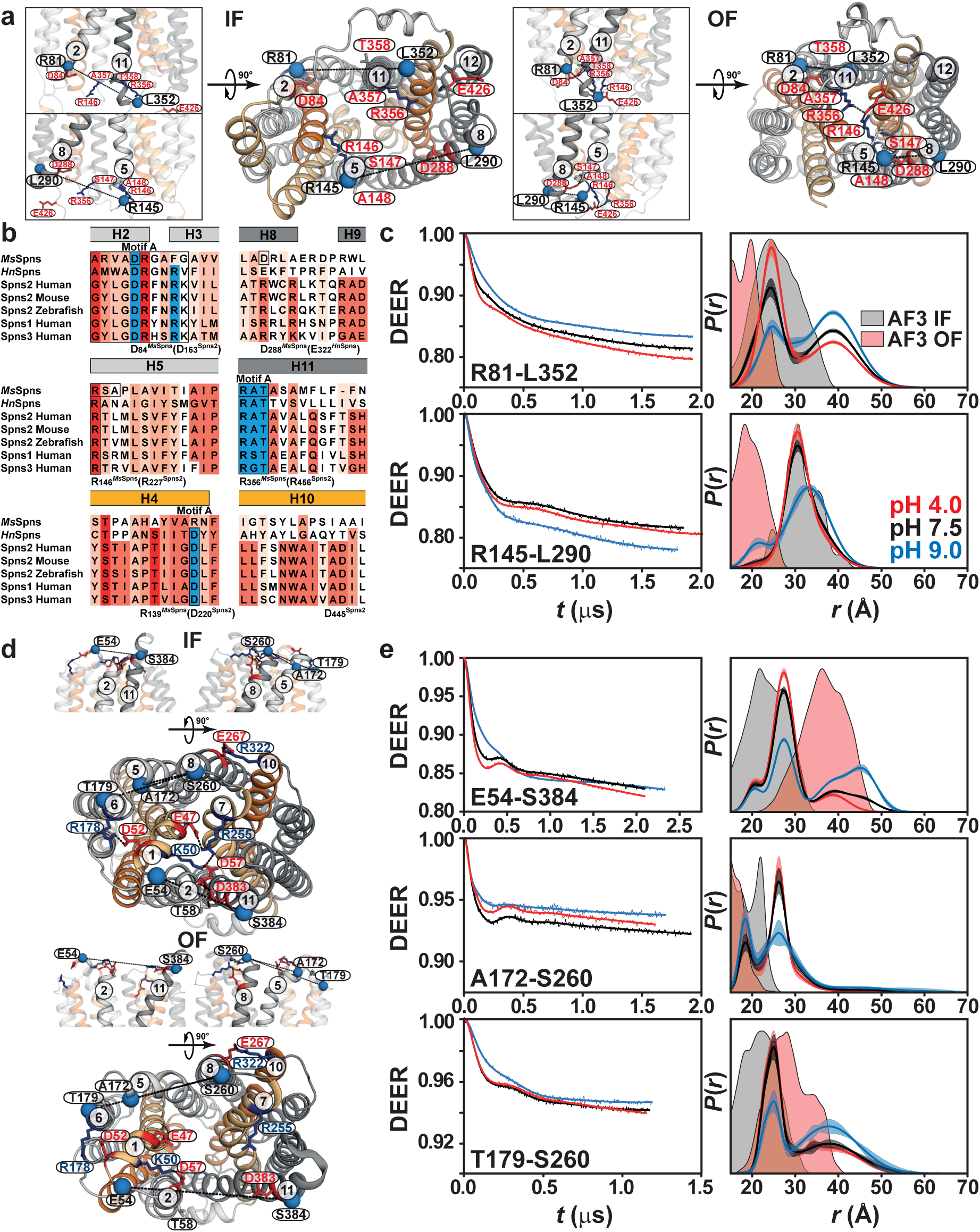
Proton binding opens the intracellular gates and closes the extracellular gates of *Ms*Spns. (**a**) Spin-label pairs (blue spheres) used for DEER distance measurements across intracellular gating helices in the NTD and CTD of IF and OF *Ms*Spns models. Membrane-view insets show the corresponding gate architecture. Functional residue interactions that stabilize the OF conformation are highlighted in red. (**b**) Sequence alignment of Spns proteins and bacterial homologs, with residues shaded by conservation in red. Residues forming structural motif A are shown in blue, and *Ms*Spns residues that stabilize the OF state are boxed. (**c**) Raw DEER decays with fits (left) and the corresponding distance dis1tributions, *P*(*r*) (right), measured in nanodiscs for R81–L352 and R145–L290 under acidic, neutral and basic pH conditions. Confidence bands (2σ) are shown about the best fit lines. Confidence bands represent uncertainty in *P*(*r*) associated with fitting of the primary DEER traces. Distance distributions predicted from the IF and OF models are shaded gray and red, respectively. (**d**) Membrane views showing the architecture of the extracellular gates. Periplasmic views show the proton-sensing network as sticks, with DEER spin-label pairs indicated as blue spheres. (**e**) DEER decays with fits and corresponding distance dis1tributions for E54-S384 at the extracellular TM2/TM11 gate, and for A172-S260 and T179-S260 at the TM5/TM8 gate.

Upon *Ms*Spns deprotonation at pH 9, the intracellular TM2/TM11 and TM5/TM8 gates become highly dynamic (Fig. 2c). Deprotonation shifts the two gate ensembles in opposite directions, favoring a closed state for TM5/TM8 and an open state for TM2/TM11, although a closed TM2/TM11 population remains detectable at pH 7.5. Protonation at pH 4 reverses these shifts. Importantly, protonation does not close the TM2/TM11 gate as predicted by the OF model; instead, it favors an IF arrangement and promotes opening of the intracellular side.

The proton-dependent distance changes observed for reporter pairs on the cavity helices (Figs. 3a and 3b) closely mirror those of the gating helices. The dominant deprotonated and protonated intermediates match the distances predicted by the OF and IF models, respectively, supporting a protonation-driven shift toward intracellular opening. By contrast, the W22-N140 pair, which reports on TM1-TM4 intradomain flexibility, shows no detectable protonation-dependent change (Supplementary Fig. 3b), consistent with a predicted Asp11^TM1^-Arg139^TM4^ salt bridge that may constrain local TM1-TM4 motion.

**Figure 3.**
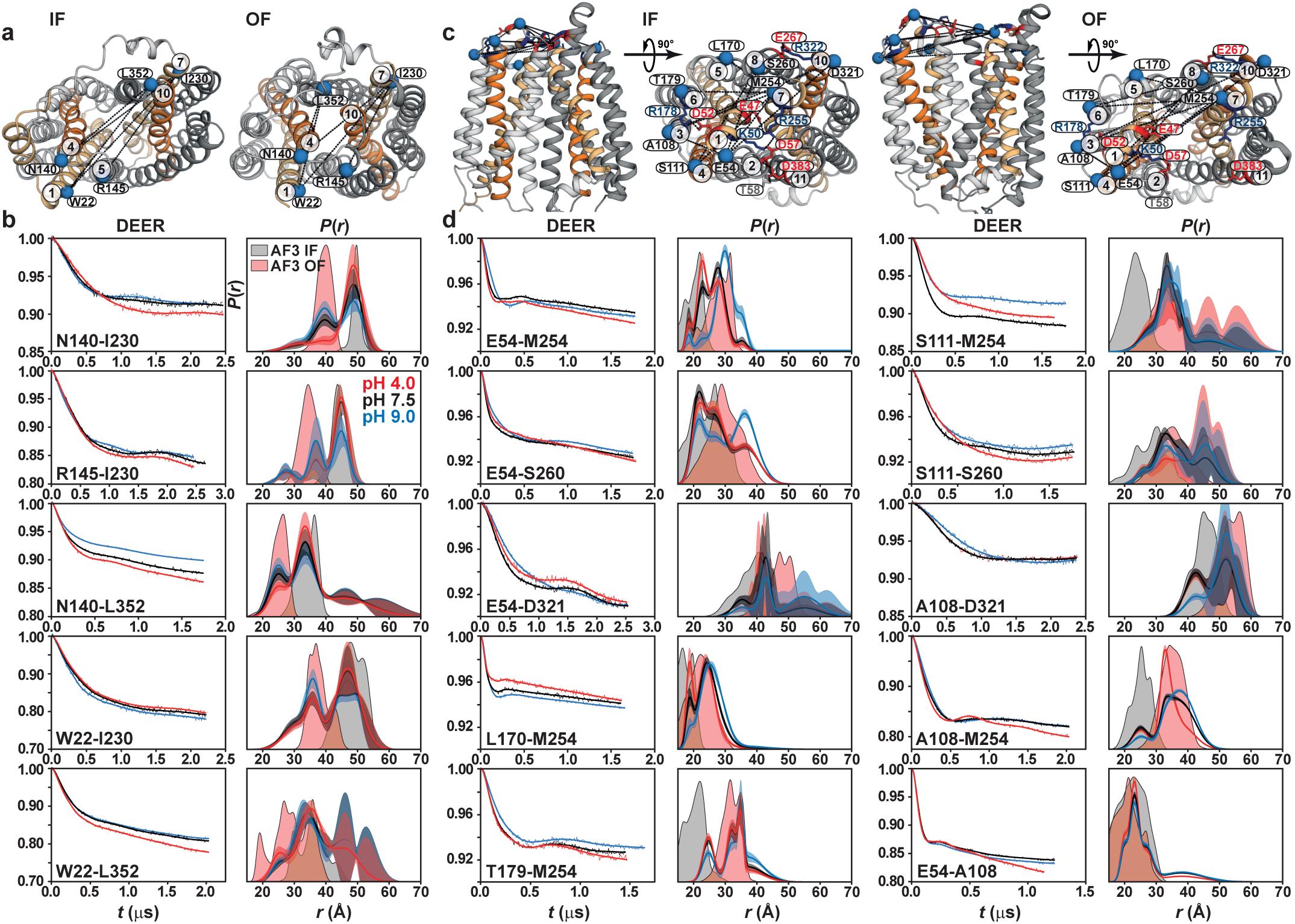
Protonation opens the intracellular substrate-binding cavity and remodels the proton-sensing network to close the extracellular side. (**a**) Spin-label pairs (blue spheres) monitoring cavity-associated helices are shown on the intracellular side of the IF and OF models. (**b**) Raw DEER decays with fits (left), together with the corresponding distance distributions, *P*(*r*) (right), measured in nanodiscs for the indicated spin-label pairs at acidic, neutral and basic pH conditions. (**c**) Spin-label pairs reporting on the extracellular proton-sensing network and binding cavity are shown on the IF and OF models. (**d**) DEER decays with fits and corresponding distance distributions, *P*(*r*), for extracellular proton-sensing and cavity-associated spin-label pairs. Distance distributions predicted from the IF and OF models are shown as gray and red shaded regions, respectively.

Distance pairs involving the support helices TMs 3, 6, 9, and 12 (Figs. 4a and 4b) follow the same global proton-dependent transition observed for the gating helices. In the deprotonated state, these support helices sample broader conformational ensembles, whereas protonation shifts the equilibria toward shorter, IF-like distances. For the A90–L298 and L197–L298 distance pairs (TM3/6–TM9), the OF state is only marginally populated under deprotonated conditions. By contrast, intradomain support-helix pairs (A90-L197, I230-L298, I230-R421, W22-A90, W22-L197, and L298-L352; Fig. 4 and Supplementary Fig. 3) show minimal protonation-dependent changes. Two conserved CTD intradomain interactions in mycobacterial Spns proteins^15^, a salt bridge between Asp294^TM9^ and Arg421^TM12^ and a hydrogen bond between Ser227^TM7^ and Asp423^TM12^ (Fig. 4a), remain intact throughout these transitions, as indicated by the narrow, proton-independent distance distributions of the corresponding reporter pairs (Fig. 4c). This suggests that alternating access in *Ms*Spns is driven primarily by interdomain gating and cavity rearrangements superimposed on a relatively stable intradomain scaffold (Figs. 2–4 and Supplementary Fig. 3).

**Figure 4.**
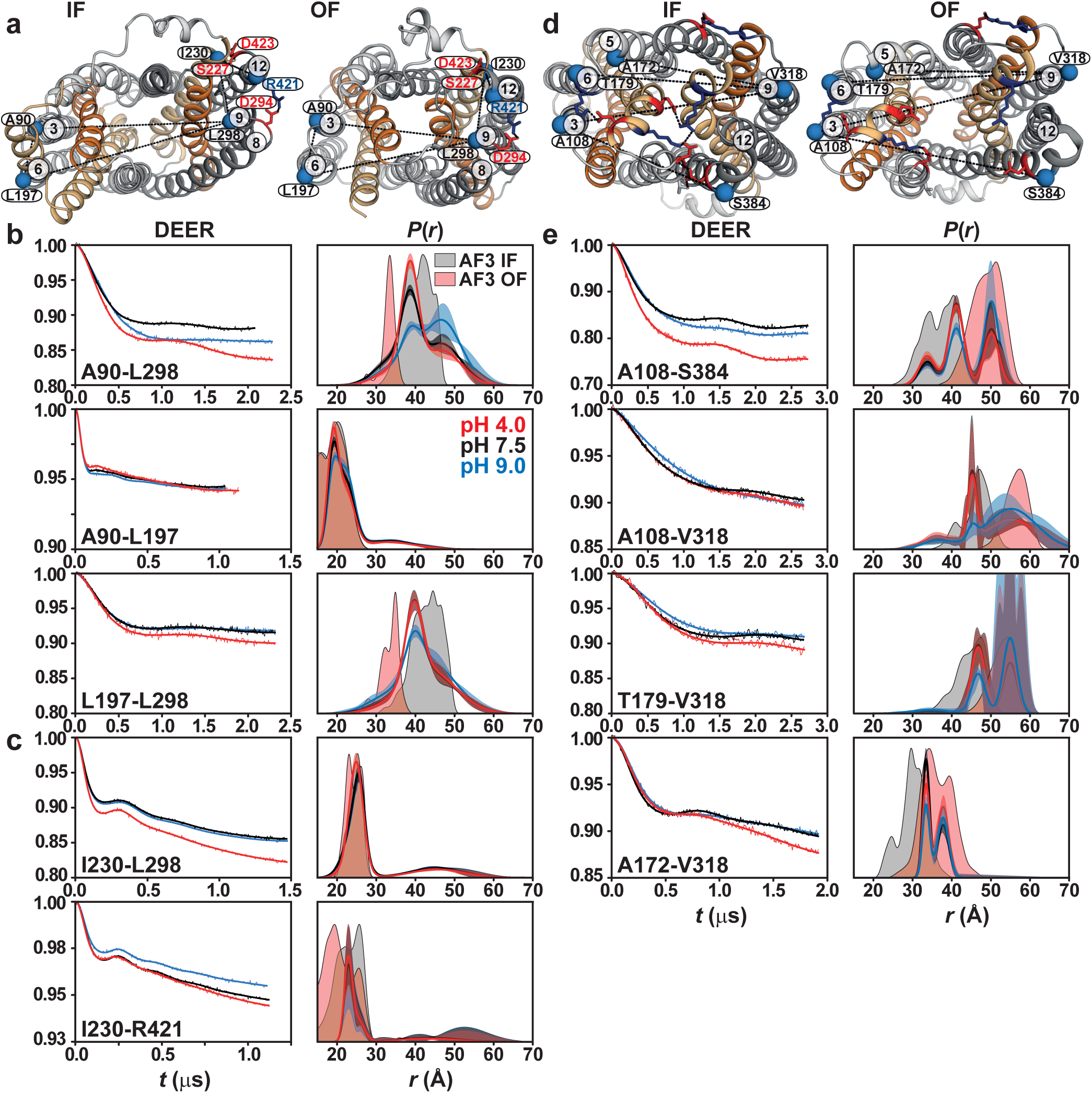
Protonation rearranges support helices preserving an IF state in *Ms*Spns. (**a**) Spin-label pairs reporting on the support helices are shown as blue spheres on the intracellular side of the IF and OF models. Intradomain interacting residues are shown as sticks. (**b**) Raw DEER decays with fits (left) and corresponding distance distributions, *P*(*r*) (right), for intracellular support-helix spin-label pairs. (**c**) Intracellular distance pairs reporting on intradomain salt-bridge and hydrogen-bond interactions. (**d**) Spin-label pairs reporting on the extracellular regions of the support helices. (**e**) DEER decays with fits and corresponding distance distributions, for extracellular support-helix spin-label pairs. Distance distributions predicted from the IF and OF models are shown as gray and red shaded regions, respectively.

### Protonation promotes closure of the extracellular side

On the extracellular, cell-envelope side, the salt bridge between an aspartate on TM2 and an arginine on TM7 (Asp57^TM2^:Arg255^TM7^ in *Ms*Spns; Fig. 1a) is among the most conserved signature interactions in bacterial and mammalian Spns transporters. Together with additional interacting residues, including a highly conserved aspartate on TM11, this interface forms an extracellular proton-sensing network proposed to drive the IF-to-OF transition (Figs. 1a and 2d). Variation in the participating residues and interaction types likely contributes to mechanistic diversity across the family^10^. For example, in human Spns3, which lacks the TM7 arginine, the equivalent salt bridge may instead be formed by the preceding lysine residue (Fig. 1a).

Under acidic conditions, protonation closes the extracellular TM2/TM11 and TM5/TM8 gating pairs (Fig. 2e). As on the intracellular side, the deprotonated state is most consistent with a broad OF-like ensemble, indicating disruption of the proton-sensing network. Notably, for the TM5/TM8 distance pair A172-M260, the average distance is shorter in the deprotonated state despite the OF conformation. In addition, unlike the pronounced opening of TM2/TM11, opening of the TM5/TM8 gate is modest during the IF-to-OF transition (Figs. 2d and 2e).

Reporter pairs spanning the cavity helices and proton-sensing network show the same proton-dependent remodeling (Figs. 3c and 3d). The E54-M254 pair, which directly monitors the highly conserved TM2-TM7 salt bridge, shows that deprotonation disrupts this interaction and stabilizes the OF state. Additional pairs spanning TM2/TM8, TM5/TM7, and TM6/TM7 support this transition, revealing an extensive extracellular proton-sensing network across TM1/TM2, TM5/TM6, TM7/TM8, and TM11. For pairs linking TM4 with TM7/8 cavity helices (S111-M254/S260; Fig. 3d), protonation stabilizes a long-distance intermediate that is absent from the IF state and only weakly sampled in the OF state. Nevertheless, consistent with the overall trend, the OF state remains dominant under deprotonated conditions for these pairs. As predicted by the models, the intradomain E54-A108 pair shows only marginal changes.

Distance pairs involving extracellular support helices likewise follow the global protonation-dependent OF-to-IF transition (Figs. 4d and 4e).

### Conserved proton switches differentially shape global and local conformational states

Spns transporters use membrane-embedded protonation switches to couple proton binding, conformational transitions and substrate transport^7^. We previously showed that, in *Hn*Spns, local electrostatic interactions among conserved membrane-embedded residues, Asp41^TM1^, Arg122^TM4^, Glu129^TM4^ and the putative substrate-binding residue Arg42^TM1^, regulate Glu129 protonation and thereby trigger larger-scale transitions, including closure of the intracellular TM2/TM11 gate. Consistent with this model, DEER measurements of protonation-mimetic mutants showed that Asp41 and Glu129 have opposing effects on intracellular conformational dynamics, while the periplasmic side is comparatively less affected, indicating weak transmembrane coupling in *Hn*Spns. Conservation of this network across Spns proteins (Fig. 1), together with impaired or reduced S1P export by the corresponding Spns2 mutants R200S^3,4,6^, D118A and E207A/E207Q^3^, and impaired LPC uptake by Spns1 R76A and E164A/E164K mutants^8,10^, suggests that analogous proton-coupling principles extend to mammalian Spns transporters.

To directly identify the protonation switches responsible for the pH-dependent conformational changes in *Ms*Spns, we systematically introduced protonation-mimetic substitutions into DEER distance pairs. The substitutions were designed to neutralize conserved acidic residues positioned within the membrane-embedded protonation-switch network or extracellular proton-sensing pathway: E126Q on TM4, D38N on TM1, the combined D38N-E126Q mutant, and D57N on TM2 (Figs. 5-7 and Supplementary Figs. 4-7).

**Figure 5.**
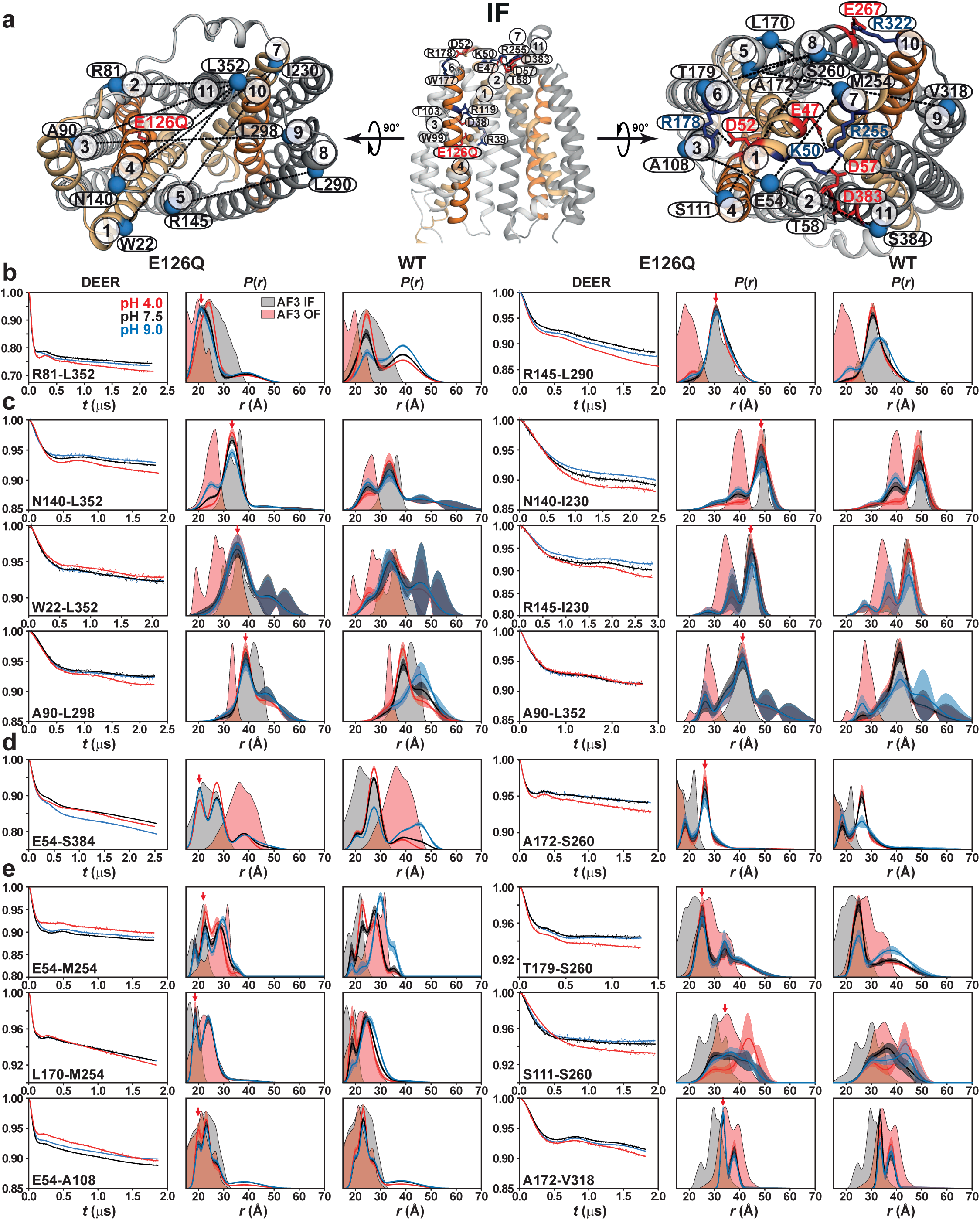
Protonation of Glu126 stabilizes the inward-facing conformation of *Ms*Spns. (**a**) Spin-label pairs reporting on intracellular and extracellular conformational changes are shown as blue spheres on the IF model. (**b**-**e**) Effects of the protonation-mimetic E1260 mutation on conformational equilibria at the intracellular gates (**b**), intracellular cavity and support helices (**c**), extracellular gates (**d**), and extracellular cavity and support helices (**e**) (Supplementary Figs. 4 and 5). The E1260 mutation was combined with the double-cysteine mutations. DEER measurements reveal that Glu126 protonation stabilizes an IF conformation, supporting its role as a protonation switch. Confidence bands (2σ), which represent the estimated uncertainty in *P*(*r*), are shown about the best fit lines. Red arrows indicate the stabilized conformational state. Glu126 protonation closes the intracellular TM2/TM11 gate while opening the TM5/TM8 gate and expanding the substrate-binding cavity to enable substrate entry.

Neutralization of Glu126^TM4^ by E126Q stabilizes an IF-like conformation (Fig. 5 and Supplementary Figs. 4 and 5). Intracellularly, E126Q closes the TM2/TM11 gate while opening the TM5/TM8 gate (Fig. 5b) and substrate-binding cavity (Fig. 5c), consistent with a substrate-entry-competent IF state. Extracellularly, the same mutation closes the TM2/TM11 and TM5/TM8 gates (Fig. 5d) and narrows the extracellular cavity and support-helix reporters (Fig. 5e). Thus, Glu126 acts as a major protonation switch: its protonated/neutral state favors inward accessibility and extracellular closure, whereas its deprotonated state contributes to the broader OF-like ensemble observed at basic pH. Notably, the extracellular response to Glu126 protonation is more heterogeneous than the intracellular response, suggesting that Glu126 may not act as the dominant protonation switch on the extracellular side.

By contrast, Asp38^TM1^ neutralization by D38N shifts *Ms*Spns toward OF-like conformations (Fig. 6 and Supplementary Fig. 6). Intracellularly, D38N closes the TM5/TM8 gate and substrate-binding cavity while opening the TM2/TM11 gate (Fig. 6c), opposing the effects of E126Q for several reporters. Extracellularly, D38N stabilizes a more open substrate-binding cavity (Fig. 6b and Supplementary Fig. 6c). However, for the extracellular TM2/TM11 and TM5/TM8 gates, monitored by E54-S384 and A172-S260, respectively, Asp38 protonation produces a more open state than Glu126 protonation but does not fully shift these reporters to the OF conformation (Fig. 6b). This suggests that complete extracellular opening requires additional elements of the extracellular proton-sensing network. Together, these data identify Asp38 as a protonation switch that favors OF-like states when neutralized.

**Figure 6.**
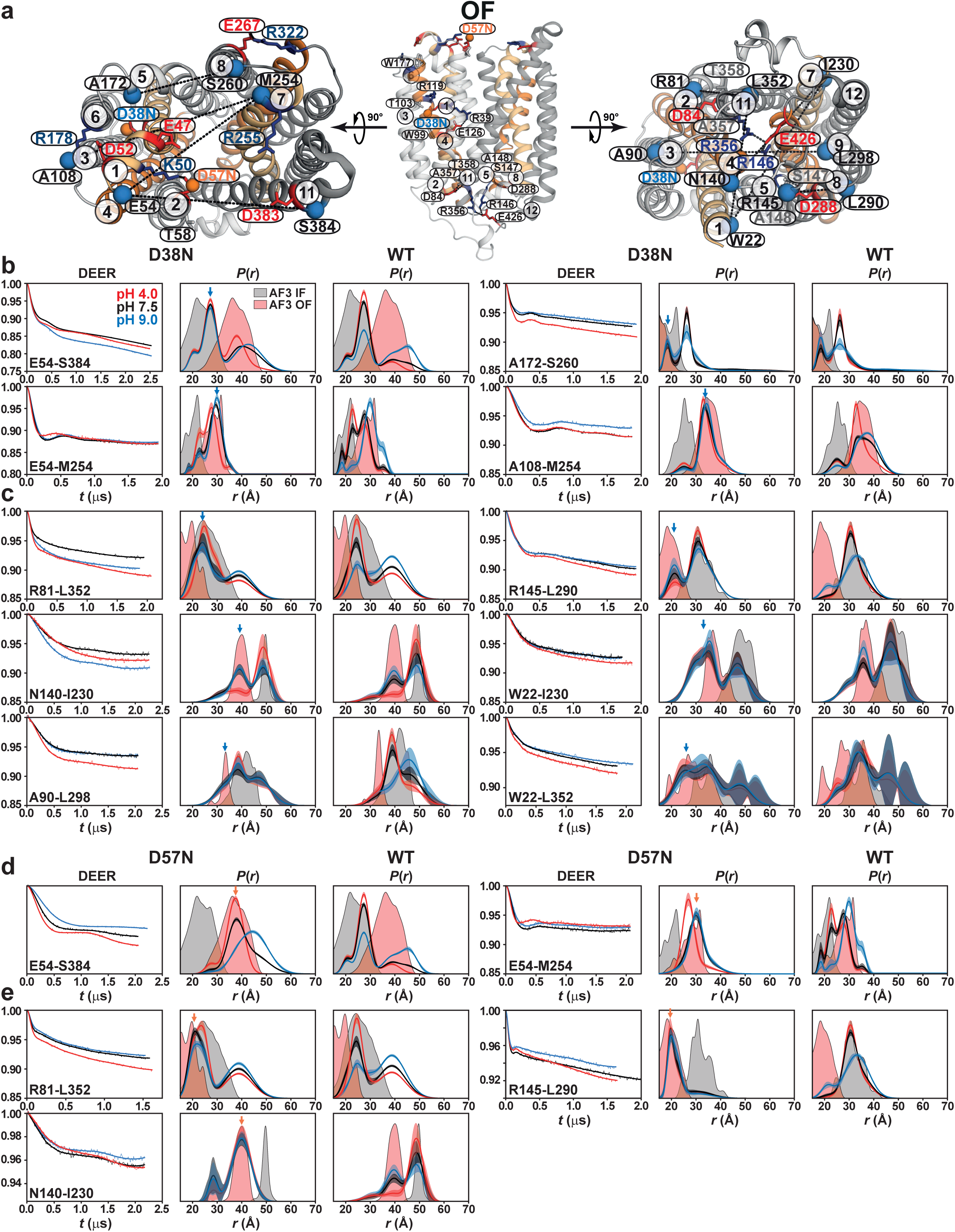
Protonation of Asp38 and Asp57 stabilizes outward-facing conformations of *Ms*Spns. (**a**) Spin-label pairs reporting on intracellular and extracellular conformational changes are shown as blue spheres on the OF model. (**b**-**e**) Effects of the protonation-mimetic D38N (Supplementary Fig. 6) and D57N mutations on conformational equilibria on the extracellular (**b,d**) and intracellular (**c,e**) sides. DEER measurements show that Asp38 protonation stabilizes an OF conformation, which is further reinforced by Asp57 protonation, supporting their roles as protonation switches and sensors. Blue and orange arrows indicate the stabilized conformational states. Asp38 protonation closes the intracellular TM5/TM8 gate and substrate-binding cavity, consistent with stabilization of an OF state.

**Figure 7.**
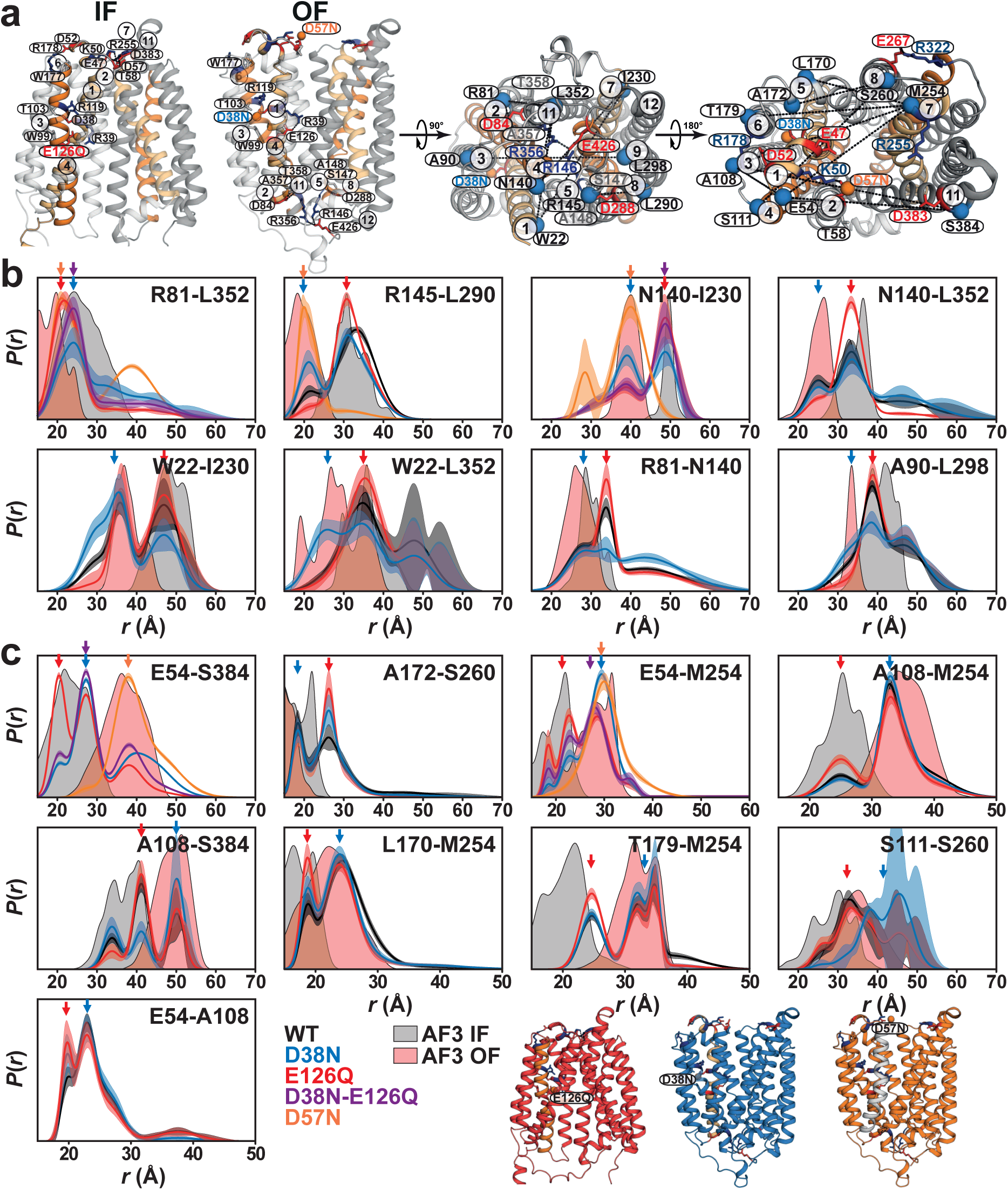
Opposing protonation switches differentially control intracellular and extracellular *Ms*Spns conformations. (**a**) Membrane views of the IF and OF models showing the membrane-embedded protonation switches and proton-sensing networks on both sides of the membrane. Spin-label pairs reporting on intracellular and extracellular conformational changes are shown as blue spheres on the OF model. (**b**) Overlay of DEER-derived distance distributions, *P*(*r*), for WT and the protonation-mimetic mutants D38N, E126Q, D38N-E126Q (Supplementary Fig. 7), and D57N on the intracellular side. E126Q shifts the intracellular gates and substrate-binding cavity toward an IF-like conformation, whereas D38N has the opposite effect and promotes OF-like closure of the intracellular side. (**c**) Corresponding extracellular-side distance distributions showing dominant control by D38N, which stabilizes OF-like extracellular opening; this OF bias is further reinforced by D57N. The D38N–E126Q double mutation underscores the opposing effects of Asp38 and Glu126 protonation and their region-specific contributions to intracellular and extracellular conformational equilibria. Colored arrows mark mutation-stabilized conformational states.

Importantly, neutralization of Asp57, the most conserved Spns signature residue (Fig. 1a), by D57N globally shifts the equilibrium toward the OF conformation, with effects that propagate from the extracellular to the intracellular side (Figs. 6a, 6d and 6e). Thus, Asp57 functions as an extracellular proton-sensing residue coupled to the global proton-coupling network. Consistent with this conserved role, recent mutational studies of human Spns2 show that the corresponding Asp137 contributes to transport activity: D137A substantially reduces S1P export, and disruption of the D137:R342 pair (D57:R255 in *Ms*Spns) impairs FTY720-P transport^3,4^. Similarly, recent studies of human Spns1 identify D94, corresponding to Spns2 Asp137 and *Ms*Spns Asp57, as a central component of the lysosomal luminal proton-sensing network; D94A markedly reduces LPC transport, whereas D94N partially rescues activity, supporting a role for protonation-mediated neutralization of this acidic residue during transport^10^.

The D38N-E126Q double mutation reveals the balance between these opposing proton switches (Fig. 7 and Supplementary Fig. 7). Rather than trapping *Ms*Spns in a single canonical IF or OF state, combined protonation of Asp38 and Glu126 enriches an IF-like intermediate while retaining OF-like subpopulations, indicating that these residues are not redundant protonation sites. Instead, they exert opposing, region-specific effects on intracellular and extracellular conformational equilibria. Asp38 is the dominant switch stabilizing the open TM2/TM11 gate on both sides of the membrane, although full extracellular TM2/TM11 opening requires Asp57 protonation (R81-L352 and E54-S384; Figs. 7b and 7c). By contrast, Asp38 protonation shifts the intracellular TM5/TM8 gate toward a closed conformation. Thus, Glu126 and Asp38 exert opposing effects on TM5/TM8 gating on both sides of the membrane, differentially stabilizing open and closed states of the intracellular and extracellular reporters (R145-L290 and A172-S260; Figs. 7b and 7c). Similarly, at the substrate-binding cavity and support helices, Glu126 protonation favors intracellular opening and extracellular closure, whereas Asp38 protonation produces the opposite effect. Thus, Glu126 primarily controls the intracellular binding cavity, while Asp38 predominantly regulates extracellular-side opening (Fig. 7). Together, the protonation-mimetic data identify Glu126 as the principal switch favoring inward accessibility, Asp38 as a switch favoring the outward-facing state, and Asp57 as an extracellular sensor that reinforces the OF conformation.

### Substrate-binding cavity of *Ms*Spns senses the proton differently

We next examined the pH-dependent conformational equilibrium of ligand-free *Ms*Spns in lipid nanodiscs using intracellular and extracellular reporter pairs, with and without protonation-mimetic substitutions (Fig. 8 and Supplementary Figs. 8 and 9). Changes in the population of increasing-distance states as a function of pH were used to estimate apparent pK values for conformational transitions (Fig. 8). In WT *Ms*Spns, nonlinear least-squares fitting yielded basic pK values of 7.7± 0.0 (*n* ≈ 2.5±0.7) and 8.0±0.1 (*n* ≈ 1.7±0.7) for the intracellular pairs A90^TM3^-L298^TM9^ and R145^TM5^-L290^TM8^, which report on support-and gating-helix motions, respectively (Figs. 8a and 8b). Because this apparent pK reflects protonation/deprotonation of acidic residue(s) that drive *Ms*Spns isomerization, the elevated values likely arise from buried sites in the central cavity, and from motif A-associated hydrogen-bonding interactions between Asp84^TM2^ and Ala357/Thr358^TM11^ and between Asp288^TM8^ and Ala148/Ser147^TM5^ (Fig. 7a). These interactions are expected to be stronger when the aspartates are deprotonated and act as hydrogen-bond acceptors, consistent with deprotonation favoring intracellular closure. The steep apparent Hill coefficients indicate a highly cooperative, switch-like transition, suggesting that multiple protonation/deprotonation events are energetically coupled into a concerted macroscopic response rather than acting independently.

**Figure 8.**
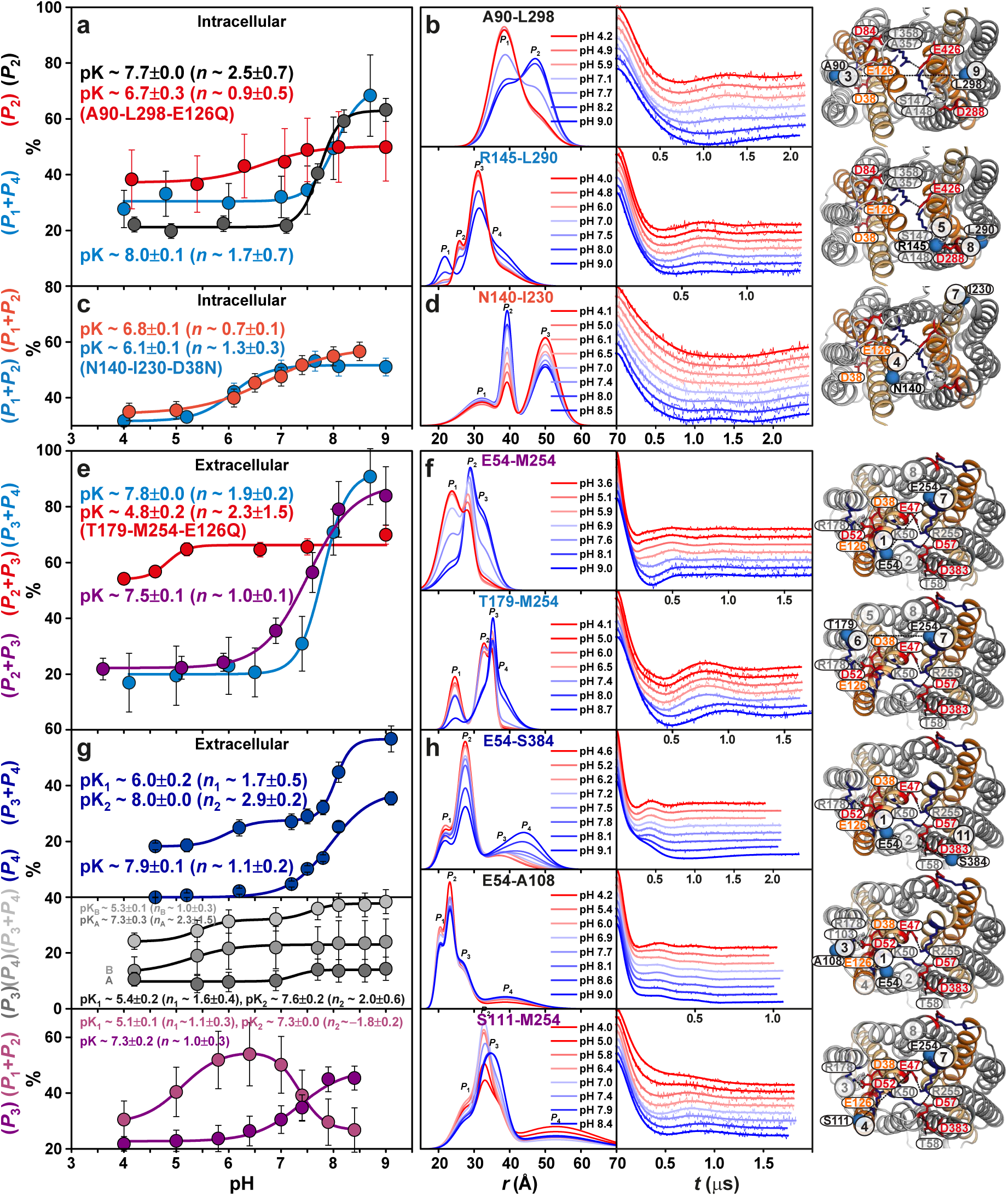
Distinct proton sensitivity and cooperativity within the *Ms*5pns substrate-binding cavity. (**a,b**) Baseline-corrected and normalized DEER traces with fits, corresponding distance dis1tributions, and pH-dependent population changes for the intracellular support-helix pair A90–L298, with and without the protonation-mimetic E126Q mutation (Supplementary Fig. 8), and for the intracellular gating pair R145–L290. Population changes in the increasing-distance peaks were used to estimate apparent pK values for conformational transitions on each side of the membrane. Glu126 protonation strongly regulates intracellular conformational dynamics. (**c,d**) Intracellular substrate-binding cavity pair N140–I230, with and without the D38N mutation, showing that the intracellular substrate-binding cavity is only modestly affected by Asp38 protonation (Supplementary Fig. 8). (**e,f**) Extracellular substrate-binding cavity pair E54–M254, which directly reports on the conserved D57:R255 salt bridge that maintains extracellular closure, and of the extracellular pair T179–M254, which indirectly reports on the TM5/8 lateral gate, with and without the E126Q mutation (Supplementary Fig. 9). As on the intracellular side, the extracellular gate and support helices are strongly modulated by Glu126 protonation. (**g,h**) Extracellular TM2/11 gating pair E54–S384, the TM1/3 pair E54–A108, which probes opening of the putative periplasmic proton-translocation pathway along TMs 1, 3 and 6, and the TM4/7 cavity pair S111–M254. Error bars in **a, c, e** and **g** represent 2σ (95%) confidence intervals for the fitted peak populations shown in **b, d, f** and **h**, respectively.

By contrast, the intracellular N140^TM4^-I230^TM7^ pair, which reports on relative motion of the substrate-binding cavity helices, yielded a lower pK of 6.8±0.1 (*n* ≈ 0.7±0.1) (Figs. 8c and 8d). This more acidic pK likely reflects the lower proton affinity of the less buried putative protonation switch Glu126^TM4^ (Fig. 7a). A Hill coefficient near 1 indicates that the conformational transition of the cavity helices is weakly cooperative and may arise from more independent protonation events.

Introducing E126Q markedly attenuates the pH-dependent rearrangement of the intracellular support-helix pair A90-L298 and the N140-I230 substrate-cavity reporter (Fig. 8a and Supplementary Fig. 8). This result confirms that Glu126 is a dominant determinant of intracellular pH sensing and cavity remodeling. By contrast, D38N only modestly affects the intracellular substrate-binding cavity (Fig. 8c and Supplementary Figs. 8c and 8d), consistent with the Asp38 predominantly regulating extracellular-side.

On the extracellular side, the T179^TM6^-M254^TM7^ pair shows a similarly basic apparent pK of 7.8±0.0 and cooperativity of 1.9±0.2, closely matching the intracellular gating/support helices (Fig. 8e and 8f). This behavior is consistent with the structural connectivity of TM5/TM6 and TM7/TM8 at the extracellular gate. E126Q strongly modulates this extracellular reporter, indicating that Glu126 protonation is transmitted across the transporter to the TM5/TM8 lateral gate and adjacent support helices (Supplementary Fig. 9). Likewise, the E54^TM1/2^-M254^TM7/8^ cavity-helix pair, which monitors the conserved Asp57^TM2^:Arg255^TM7^ salt bridge, exhibits an elevated pK of 7.5±0.1 because it is coupled to the gating helices. Although, its lower cooperativity suggests a more independent response of the substrate-binding and release helices.

Interestingly, the extracellular pairs E54^TM1/2^-S384^TM11^, E54^TM1/2^-A108^TM3^, and S111^TM4/3^-M254^TM7/8^ display biphasic pH titration behavior, consistent with sequential protonation-linked conformational transitions (Fig. 8g and 8h). This behavior is plausible given the extensive extracellular proton-sensing network and its coupling to membrane-embedded protonation switches (Fig. 7a). For E54-S384, which directly reports on TM2/TM11 gating and is sensitive to the hydrogen-bonding interaction between Asp383^TM11^ and Asp57/Thr58^TM2^, a basic pK of 7.9±0.1 (*n* ≈ 1.1±0.2) is observed for the state corresponding to complete gate disruption (Fig. 8g), consistent with the global gating transition. For the extracellular S111-M254 cavity pair, the corresponding state exhibits a similar apparent pK and cooperativity to the E54-M254 cavity reporter, with a pK of 7.3±0.2 (*n* ≈ 1.0±0.3).

The E54-A108 pair monitors the proposed proton-transfer pathway to Asp38^TM1^ through highly conserved Spns residues (Figs. 1b, 1e, and 7a), including Trp99 and Thr103 on TM3 and Trp177 on TM6^7^. We previously proposed a conserved Spns proton-coupling mechanism in which protonation and deprotonation of the TM4 glutamate are regulated by the protonation state of the TM1 aspartate and by substrate binding^7^. Our extensive MD simulations further implicated neutralization of the highly conserved TM4 and TM1 arginine residues in this regulatory mechanism (Figs. 1 and 7a)^7^. Consistent with this model, the AlphaFold3-generated IF structure shows deprotonated Asp38^TM1^ forming a salt bridge with Arg119^TM4^ and a hydrogen bond with Thr103^TM3^ (Fig. 7a, IF). This arrangement is expected to favor protonation of Glu126^TM4^ and disruption of its salt bridge with Arg39^TM1^. In the OF model, by contrast, the Asp38^TM1^ interactions are disrupted (Fig. 7a, OF), potentially mimicking Asp38 protonation. Thus, Asp38-linked conformational changes can be transmitted directly to the extensive extracellular proton-sensing network through interactions involving Glu47, Lys50, Asp52, Asp57, and Thr58 (Fig. 8b, inset). Accordingly, for E54-A108 and A108-M254 pairs, D38N strongly suppresses pH-dependent conformational changes (Supplementary Fig. 9), supporting a dominant role for Asp38 in regulating this pathway.

Overall, the gating/support-helix network and substrate-binding cavity display distinct proton affinities, cooperativities, and sensitivities to individual protonation switches.

### Substrate hydrophobicity drives opposing shifts in *Ms*Spns conformational equilibria

Deletion of *Ms*Spns in *Mycobacterium smegmatis* increases intracellular accumulation of, and sensitivity to, water-soluble compounds such as capreomycin and ethidium bromide, while reducing susceptibility to lipophilic drugs such as rifampicin^7,15^. Consistent with an efflux antiporter mechanism, the cationic compounds capreomycin and ethidium bromide stabilize the OF conformation at pH 7.5 relative to the apo state (Fig. 9 and Supplementary Figs. 10 and 11). These OF shifts are observed on both extracellular and intracellular reporters and persist in selected protonation-mimetic backgrounds. Docking capreomycin into the IF conformation highlights highly conserved aromatic residues across the binding cavity of mycobacterial Spns proteins that may contribute to substrate binding (Fig. 9a). By contrast, lipophilic compounds, including rifampicin, shift the conformational equilibrium toward the IF state, consistent with possible uptake. Previous studies showed that the lipophilic Spns2 inhibitors 16d (SLF1081851) and 33p (SLB1122168) attenuate transport by trapping Spns2 in an inward-facing state^2,6^.

**Figure 9.**
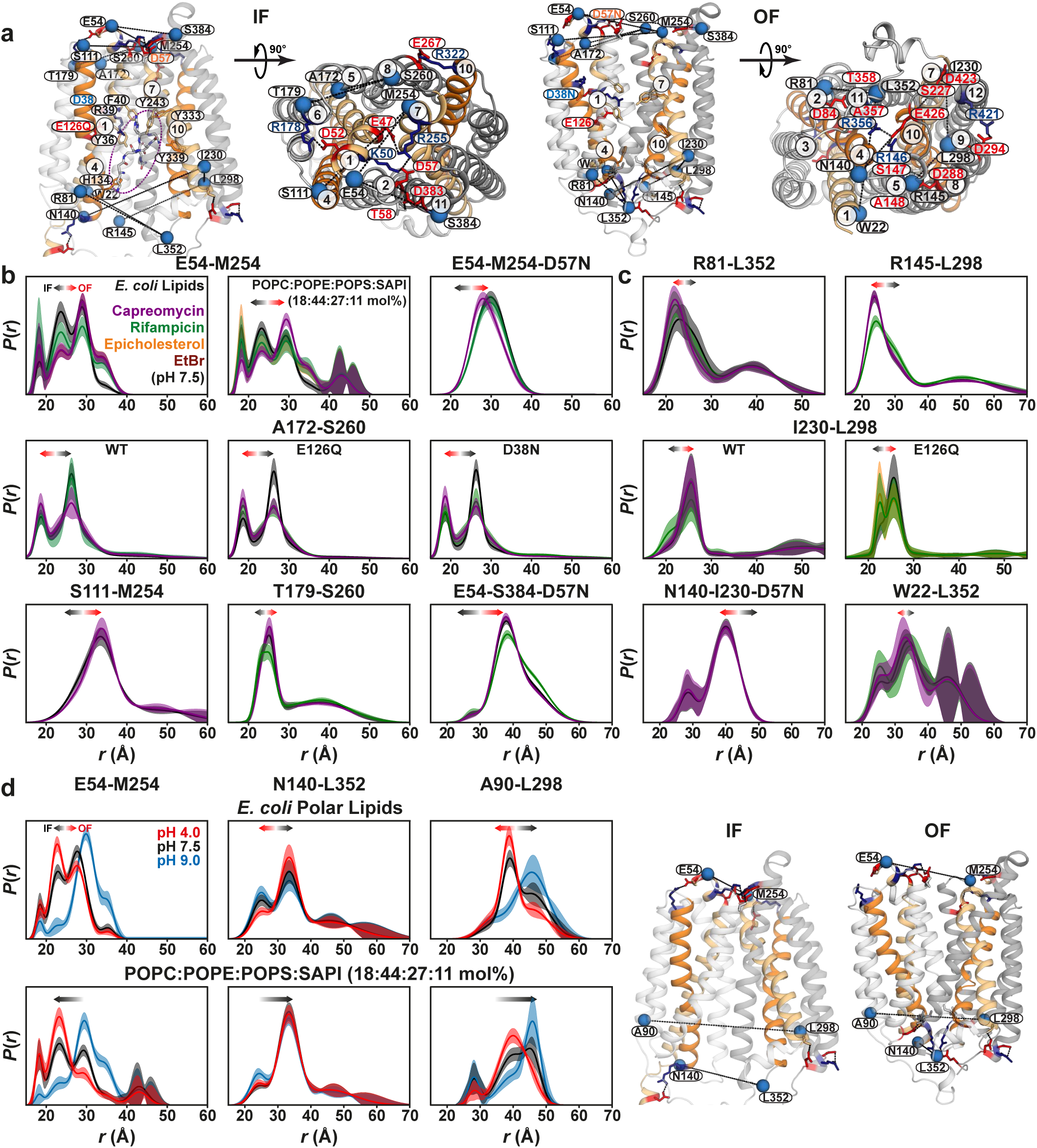
Hydrophobic and hydrophilic substrates differentially shift *Ms*5pns conformational equilibria. (**a**) Membrane and periplasmic views of the IF model and membrane and cytoplasmic views of the OF model. Capreomycin (circled) is docked in the substrate-binding cavity of the IF state, with conserved aromatic cavity residues shown as sticks. DEER spin-label pairs are shown as blue spheres. (**b,c**) DEER distance distributions measured at pH 7.5 in the apo state and in the presence of substrates on the extracellular (**b**) and intracellular (**c**) sides, with and without protonation-mimetic mutations. Two lipid compositions were tested for the extracellular pair E54-M254. Water-soluble cationic substrates (capreomycin and ethidium bromide) stabilize the OF conformation, whereas lipophilic substrates (rifampicin and epicholesterol) shift the equilibrium toward the IF state. (**d**) Effects of lipid composition on *Ms*Spns conformational equilibria probed by the extracellular pair E54-M254 and the intracellular pairs N140–L352 and A90–L298. Raw DEER decays with fits are shown in Supplementary Figs. 10-12.

Loss of *Ms*Spns was previously shown to strongly upregulate the downstream gene encoding isocitrate lyase (ICL)^15^. Defective efflux may increase intracellular accumulation of xenobiotics and metabolic by-products, thereby triggering compensatory metabolic responses. In *Mycobacterium tuberculosis,* ICL is broadly induced during antibiotic tolerance and persistence^31^, suggesting that *Ms*Spns-linked changes in ICL expression may reflect a broader stress-adaptation program. The genomic linkage of a 3-α-hydroxysteroid dehydrogenase (MSMEG_3704), *Ms*Spns (MSMEG_3705), and an ICL (MSMEG_3706) suggests a functional connection to lipid or sterol catabolism, consistent with the role of ICL in buffering acetyl-and propionyl-CoA flux. Although *Ms*Spns confers resistance to cationic antibiotics such as capreomycin, its native substrates may include cationic amphipathic molecules, sterol-derived catabolites, or other amphipathic metabolites. Consistent with this possibility, cholest-5-en-3α-ol (epicholesterol), a sterol derivative and potential lipophilic substrate, stabilizes the IF conformation, like rifampicin, based on the E54-M254, R145-I230, and I230-L298-E126Q distance pairs (Fig. 9 and Supplementary Figs. 10 and 11), suggesting uptake.

This dual behavior, together with the opposing effects of water-soluble and lipophilic compounds on conformational equilibria, parallels our findings for *Hn*Spns^7^ and may represent a shared mechanistic signature of Spns proteins.

Interestingly, the lipid composition used in the *Hn*Spns study also shifts *Ms*Spns toward the IF state across pH values (Fig. 9d and Supplementary Fig. 12), while preserving the differential effects of hydrophilic and lipophilic substrates (Fig. 9 and Supplementary Figs. 10 and 11). This finding suggests that specific lipid environments can tune the conformational landscape without overriding ligand-dependent directionality. Because phosphatidylinositol (PI) binds Spns2 with relatively high affinity among non-phosphorylated glycerophospholipids, possibly reflecting a broader preference for inositol-containing or anionic lipids in Spns proteins^5^, L-α-phosphatidylinositol (SAPI) in this lipid mixture could act as a potential *Ms*Spns substrate or allosteric modulator that stabilizes the IF conformation, similar to rifampicin or epicholesterol. The IF stabilization by PI is particularly intriguing because mycobacterial membranes contain phosphatidylinositol, an essential precursor for phosphatidylinositol mannoside, lipomannan, and lipoarabinomannan biosynthesis, which shape cell-envelope architecture and host-pathogen interactions^32,33^. Accordingly, the recent observation that Spns2 exports S1P while importing glucose^14^ supports the broader possibility that Spns transporters can couple movement of chemically distinct ligands in opposite directions.

### DEER-guided refinement and modeling of IF and OF *Ms*Spns structures

To convert the experimental DEER data into structural models, we used DEERFold^24^, a fine-tuned AlphaFold2-based network^26^ that incorporates DEER distance distributions as distogram inputs to predict spin-label-constrained conformations. Single-Gaussian approximations of the if-and OF-dominant distance distributions measured at pH 4.0 and pH 9.0, respectively, were used as DEERFold restraints. The resulting IF and OF ensembles closely matched the corresponding unbiased AlphaFold3 (AF3) models (Fig. 10a)^27^. The top IF models showed an average TM-score of 0.96 and an average Cα RMSD of 1.86 Å relative to the AF3 IF model, whereas the top OF models showed an average TM-score of 0.97 and an average Cα RMSD of 1.76 Å relative to the AF3 OF model. Refinement of the unbiased AF3 models using the same DEER distance constraints produced similarly convergent structures (Fig. 10a and Supplementary Figs. 13 and 14)^25^.

**Figure 10.**
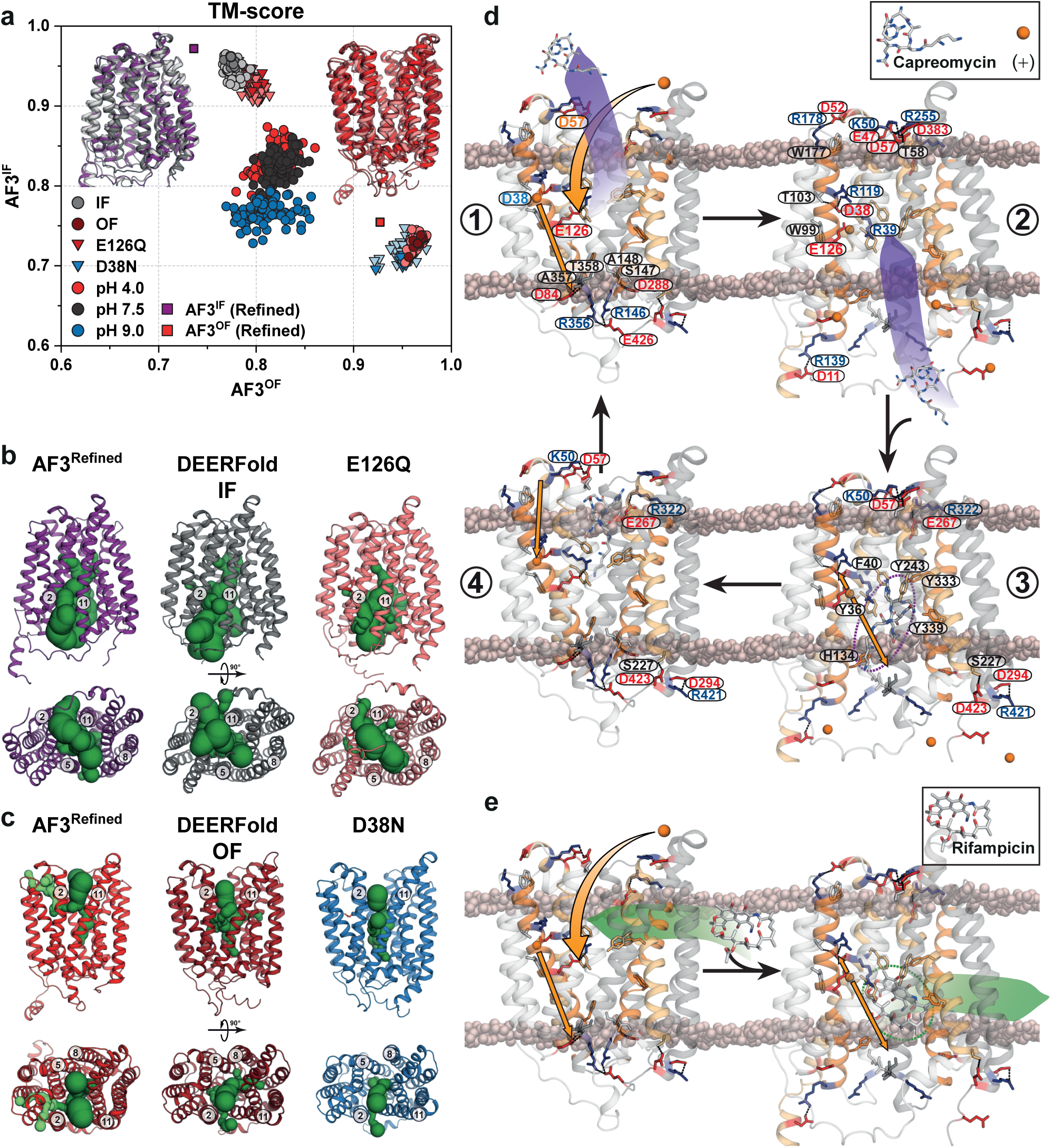
Proposed dual transport model for *Ms*Spns. (**a**) DEER-guided AlphaFold ensembles, generated by DEERFold, are compared with DEER-refined AF3 models. Models are sorted by TM-score^46^ similarity to unbiased IF and OF AF3 models, with the top 10 DEERFold IF and OF models shown in darker colors. Structural overlays of unbiased AF3 (IF: light gray, OF: light red), top DEERFold (IF: dark gray, OF: firebrick), and refined AF3 models (IF: purple, OF: red) show strong convergence. (**b**) Calculated tunnels (MOLEonline 2.5^48^, dark green) in IF models indicate that substrates can access the binding cavity from the inner membrane leaflet through the TM2/11 and TM5/8 gates or directly from the cytoplasm, allowing entry of both lipophilic and water-soluble substrates. Glu126 protonation further constricts the TM2/11 gate. (**c**) Tunnels calculated in OF models suggest that substrates access or exit the cavity from the outer membrane leaflet through the TM2/11 gate or directly from/to the periplasm. (**d**) Proposed antiport model for water-soluble cationic substrates. Asp38 protonation stabilizes the resting OF state, likely through a putative periplasmic proton tunnel involving conserved residues on TMs 3 and 6 (panel c, lime tunnel in AF3-refined model) (**1**). Glu126 protonation and proton transfer from Asp38 to intracellular acidic residues favor the higher-energy IF state and prime substrate binding (**2**). Binding of cationic substrates promotes Glu126 deprotonation and Asp38 reprotonation, disrupting the extracellular proton-sensing network and returning *Ms*Spns to the OF state (**3** to **4**). Four conserved intradomain salt bridges and hydrogen bonds remain largely intact during IF–OF isomerization (steps 3 and 4), consistent with limited proton-and substrate-dependent changes in intradomain DEER distances. (**e**) Proposed symport model for lipophilic substrates. Lipophilic substrates such as rifampicin bind from the outer membrane leaflet and, together with Glu126 protonation, drive OF-to-IF isomerization, promoting proton release to the cytoplasm and substrate release to the inner leaflet.

Using the same approach, we generated DEER-guided models of *Ms*Spns carrying the protonation-mimetic mutations E126Q and D38N, using the stabilized distance components observed under each condition. As predicted, E126Q stabilized an IF conformation closely resembling the AF3 IF model, with an average TM-score of 0.94 and an average Cα RMSD of 2.20 Å. Conversely, D38N stabilized an OF conformation with similarly high agreement to the AF3 OF model, with an average TM-score of 0.95 and an average Cα RMSD of 2.11 Å (Fig. 10a). When the complete multi-Gaussian distance distributions from all three pH conditions were used, DEERFold generated more dispersed ensembles, consistent with its fine-tuning on single-Gaussian distance distributions^24^. The best pH 4.0 model reached a TM-score of 0.86 and a Cα RMSD of 3.61 Å relative to the AF3 IF model, whereas the best pH 9.0 model reached a TM-score of 0.86 and a Cα RMSD of 3.56 Å relative to the AF3 OF model (Fig. 10a). Even with this lower-precision representation, the pH-dependent trend remained clear: protonated restraints at pH 4.0 produced more IF-like models, whereas deprotonated restraints at pH 9.0 produced more OF-like models, demonstrating that the IF-to-OF conformational shift is robust across distance-restraint representations.

Tunnel analysis of the unbiased and refined AF3 models, together with the top DEERFold models (dark green; Figs. 10b and 10c), indicates that in the IF state, the substrate-binding site is accessible laterally from the inner membrane leaflet through the TM2/TM11 and TM5/TM8 gates, as well as directly from the cytoplasm. Thus, both lipophilic and water-soluble substrates can enter or exit the transporter in this state (Fig. 10b). Glu126 protonation further constricts the TM2/TM11 gate while preserving alternative routes for substrate entry, consistent with the E126Q DEER data showing an IF-like, substrate-entry-competent ensemble (Figs. 5 and 10b). In the OF state, the extracellular TM5/TM8 membrane-facing gate is constricted, allowing substrates to access or exit the central cavity from the outer membrane leaflet through the TM2/TM11 gate or directly from the periplasm (Fig. 10c). The D38N OF model reveals similar access routes, consistent with stabilization of an OF-like conformation (Figs. 6 and 10c).

## Discussion

Using an integrated spectroscopic and computational approach, we define the proton-and substrate-coupled alternating-access mechanism of *Ms*Spns and show how a conserved Spns fold supports mechanistic versatility. DEER spectroscopy in lipid nanodiscs, combined with DEER-guided and AlphaFold-based modeling, reveals that protonation shifts *Ms*Spns toward an IF state, whereas deprotonation favors a broader OF ensemble through coordinated remodeling of the intracellular and extracellular gates. These transitions are governed by an extensive proton-sensing architecture composed of membrane-embedded protonation switches and an extracellular proton-sensing network, while the substrate-binding cavity displays distinct proton affinity and weaker cooperativity, indicating differential proton sensing within the transporter. Comprehensive protonation-mimetic mutagenesis reveals a residue-level mechanism: Glu126 protonation stabilizes an IF state competent for substrate binding through the TM5/TM8 lateral gate or directly from the cytoplasm, whereas Asp38 protonation favors an OF conformation. Subsequent Asp57 protonation further opens the extracellular side and stabilizes the transporter in the OF state. Functionally, hydrophilic cationic substrates such as capreomycin and ethidium bromide stabilize the OF state, consistent with efflux antiport, whereas lipophilic compounds such as rifampicin, epicholesterol, and selected phospholipids shift the equilibrium toward the IF state, suggesting uptake-like transport or allosteric stabilization. Together, these findings establish *Ms*Spns as a proton-coupled transporter with opposing responses to chemically distinct substrates and provide a structural framework that connects bacterial efflux and influx to the emerging mechanistic landscape of mammalian Spns proteins.

The proton-coupled ensemble shifts observed here are substantial, internally consistent across multiple reporter sites, and supported by both DEER-guided and unbiased structural models. This convergence indicates that *Ms*Spns does not toggle between two static conformations, but instead samples a proton-dependent energy landscape in which the gating helices, support helices and substrate-binding cavity are differentially coupled. Protonation-mimetic data further show that these shifts arise from specific acidic residues with distinct energetic roles. Glu126 acts as the dominant intracellular protonation switch, analogous to the TM4 glutamate in *Hn*Spns, whereas Asp38 tunes the periplasmic proton-transfer pathway and favors the OF state. Asp57, positioned within the conserved extracellular salt-bridge network, functions as an extracellular sensor that propagates protonation changes to the broader gating apparatus. Thus, protonation of Glu126, Asp38 and Asp57 is transmitted most efficiently through the interconnected gating and support-helix network, producing concerted, switch-like rearrangements of the intracellular and extracellular gates. By contrast, the substrate-binding cavity is shaped by additional local factors, including ligand chemistry and charge compensation, which partially decouple it from the global gating transition and yield lower apparent cooperativity with stronger substrate dependence.

Based on AF3 and DEER-refined IF and OF models of *Ms*Spns stabilized by protonation of conserved switches, together with its proton-and ligand-dependent conformational dynamics, we propose a dual transport model for *Ms*Spns (Figs. 10d and 10e). We previously proposed for *Hn*Spns a proton-coupling mechanism centered on highly conserved membrane-embedded protonation switches (Fig. 1b), suggesting that this mechanism is broadly conserved across the Spns family^7^. This model identifies residues critical for protonation and its regulation, explains how sequential protonation of these switches drives conformational transitions, and provides a mechanism for coupling proton movement to substrate transport, including a putative proton-translocation pathway (Fig. 10). However, stabilization of a canonical OF conformation in *Ms*Spns contrasts sharply with *Hn*Spns and instead more closely resembles the behavior of human Spns transporters, as reflected in their solved structures^2,10^. In this model, the resting state is OF (Fig. 10d, steps 1 and 4), stabilized by protonation of Asp38^TM1^, likely through a periplasmic tunnel leading to Asp38 (Fig. 10c, lime tunnel in the AF3-refined model) and involving conserved Spns residues, including Trp99 and Thr103 on TM3 and Trp177 on TM6 (Fig. 10d, step 4)^7^. The IF state (steps 2 and 3) represents a higher-energy conformation and is stabilized by protonation of Glu126^TM4^. During the OF-to-IF transition (Fig. 10d, step 1 to 2), periplasmic release of a cationic water-soluble substrate such as capreomycin primes Glu126 for protonation from the periplasmic side, while proton transfer from Asp38 to acidic residues at the intracellular gates, such as Asp84, disrupts stabilizing hydrogen bonds, favors the IF conformation^34^, and primes the transporter for substrate binding from the intracellular side (step 2). In this state, deprotonated Asp38 can form a salt bridge with Arg119^TM4^ and a hydrogen bond with Thr103^TM3^. Binding of cationic substrates then induces deprotonation of Glu126, either through proton transfer to the intracellular side or relay back to Asp38, followed by salt-bridge formation with Arg39^TM1^ (step 3). Capreomycin binding together with Asp38 protonation generates local conformational changes that propagate to the extracellular proton-sensing network, stabilizing the OF state (step 3 to 4), consistent with antiport. Notably, capreomycin, a polypeptide antibiotic, behaves as a tetraprotic acid and can also serve as a proton donor at pH 7.5^35^. Four intradomain salt bridges and hydrogen bonds (Lys50^TM1^:Asp57^TM2^, Glu267^TM8^:Arg322^TM10^, Asp294^TM9^:Arg421^TM12^, and Ser227^TM7^-Asp423^TM12^) remain intact during IF-to-OF isomerization (Fig. 10d, steps 3 and 4), consistent with the limited proton-and substrate-dependent changes in most intradomain DEER distances (Figs. 3d, 4c, and Supplementary Fig. 3b). By contrast, binding of lipophilic substrates such as rifampicin directly from the outer membrane leaflet, together with Glu126 protonation, drives OF-to-IF isomerization and release of protons to the cytoplasm and substrate to the inner leaflet, consistent with symport-like uptake (Fig. 10e).

This transport model reveals previously unrecognized features of alternating access in Spns proteins and shows how a conserved set of proton-sensing interactions can be repurposed to support multiple transport outputs (Figs. 1, 2b, 7a)^7^. It also places *Ms*Spns within a broader MFS framework that includes proton-coupled sugar acid and sugar symporters such as *E. coli* DgoT and XylE^36,37^. The proposed regulatory role of Asp38 in *Ms*Spns and Asp41 in *Hn*Spns is reminiscent of Asp27 in XylE, while the reversible protonation of Asp46 and Glu133 in DgoT parallels the membrane-embedded protonation switches of Spns proteins^7,37,38^. Together with the recent finding that Spns2 exports S1P while importing glucose^14^, our observation that SAPI potentially shifts *Ms*Spns toward the IF state (Fig. 9d) supports a flexible model in which ligands and lipids tune transport directionality by biasing the same proton-coupled conformational landscape. Alongside our findings in *Hn*Spns^7^, these results suggest that dual transport may be an emerging feature of the Spns fold, whereby one ligand gradient can promote counter-transport of another and lessen the requirement for strict proton coupling under some conditions^2,14^. More broadly, ligand chemistry, particularly the pH-dependent balance between water solubility and lipophilicity in substrates such as S1P, may determine whether a ligand is exported, imported, or acts as an allosteric modulator^2,14^.

The dual behavior of *Ms*Spns, together with the expanding functional diversity of the Spns family, indicates that the same core mechanistic elements, including membrane-embedded protonation switches and extracellular proton-sensing networks, can power substrate movement in opposite directions. The protonation-mimetic mutations reported here provide a framework for separating the energetic roles of these elements in bacterial, human, and orphan Spns transporters. In this context, the ability of lipophilic compounds to favor IF states highlights a therapeutically important possibility: such modulators may either be transported by Spns proteins^15^ or trap them in inward-facing conformations^2,6^. Applying this integrated structural-dynamics platform to additional Spns transporters should define the shared and divergent principles that underlie Spns energy coupling, ligand directionality, and regulation.

## Methods

No statistical methods were used to predetermine sample size. The experiments were not randomized. The investigators were not blinded to allocation during experiments and outcome assessment.

### Site-directed mutagenesis

Codon-optimized *Ms*Spns (GenScript) was cloned into pET19b vector encoding an N-terminal 10-His tag under control of an inducible T7 promoter. The three cysteine residues in *Ms*Spns were mutated (C95S, C325A, C412L) via site-directed mutagenesis with complementary oligonucleotide primers, yielding the CL protein. This construct was used as the template to introduce double-cysteine pairs. Protonation-mimetic mutations E126Q, D38N, D38N-E126Q, and D57N were introduced into the relevant double-cysteine backgrounds using the same site-directed mutagenesis strategy. Substitution mutations were generated using a single-step PCR in which the entire template plasmid was replicated from a single mutagenic primer. *Ms*Spns mutants were sequenced using both T7 forward and reverse primers to confirm mutagenesis and the absence of aberrant changes. Mutants are identified by the native residue and primary sequence position followed by the mutant residue.

### Expression, purification, and labeling of *Ms*Spns

*Ms*Spns was expressed and purified using a similar protocol as previously published^7^. *Escherichia coli* C43 (DE3) cells (Sigma-Aldrich) were freshly transformed with pET19b vector encoding recombinant *Ms*Spns mutants. A transformant colony was used to inoculate Luria–Bertani (LB) media (Fisher Bioreagents) containing 0.1 mg/mL ampicillin (Gold Biotechnology), which was grown overnight (∼15 h) at 34 °C and was subsequently used to inoculate 3-6 L of minimal medium A at a 1:50 dilution. Cultures were incubated while being shaken at 37 °C until they reached an absorbance at 600 nm (Abs_600nm_) of ∼ 0.8, at which time *Ms*Spns expression was induced by the addition of 1 mM IPTG (Gold Biotechnology). The cultures were incubated overnight (∼15 h) at 20 °C and then harvested by centrifugation. Cell pellets were resuspended in resuspension buffer (20 mM Tris⋅HCl, pH 8.0, 20 mM NaCl, 10 mM imidazole, and 10% [vol/vol] glycerol) at 15 mL per liter of culture, including 10 mM DTT, 1 mM EDTA, and 1 mM PMSF, and the cell suspension was lysed via sonication. Cell debris was removed by centrifugation at 9,000 × *g* for 10 min. Membranes were isolated from the supernatant by centrifugation at 180,000 × *g* for 1.5 h.

Membrane pellets were solubilized in resuspension buffer (15 mL/g membrane) containing 5 mM LMNG (Anatrace) and 0.5 mM DTT and incubated on ice with stirring for 1 h. Insoluble material was cleared by centrifugation at 180,000 × *g* for 30 min. The cleared extract was bound to 1.0 mL (bed volume) of Ni-NTA Superflow resin (Qiagen) at 4 °C for 2 h. After washing with 10 bed volumes of buffer containing 30 mM imidazole and 0.2 mM LMNG, *Ms*Spns was eluted with buffer containing 300 mM imidazole.

Double-cysteine mutants were labeled with two rounds of 20-fold molar excess 1-oxyl-2,2,5,5-tetramethylpyrroline-3-methyl methanethiosulfonate (Enzo Life Sciences) per cysteine on ice in the dark over a 4-h period, after which the sample was kept on ice at 4 °C overnight (∼15 h) to yield the spin label side chain R1. Unreacted spin label was removed by size exclusion chromatography over a Superdex200 Increase 10/300 GL column (GE Healthcare) into 50 mM Tris/MES, pH 7.5, 75 mM NaCl, 0.2 mM LMNG, and 10% (vol/vol) glycerol buffer. Peak fractions of purified *Ms*Spns were combined and the final concentration was determined by A_280_ measurement (ε = 57,410 M^−1^⋅cm^−1^) for use in subsequent studies.

### Reconstitution of *Ms*Spns into nanodiscs

For all distance pairs, *E. coli* Polar Lipid Extract (Avanti Polar Lipids) was used. For a subset of distance pairs, 1-palmitoyl-2-oleoyl-sn-glycero-3-phosphocholine (POPC), 1-palmitoyl-2-oleoyl-sn-glycero-3-phosphoethanolamine (POPE), 1-palmitoyl-2-oleoyl-sn-glycero-3-phospho-L-serine (POPS) and L-α-phosphatidylinositol (Liver, Bovine, SAPI) (Avanti Polar Lipids) were combined in a 17.5:44:27.5:11 (mol/mol) ratio, dissolved in chloroform, evaporated to dryness on a rotary evaporator, and desiccated overnight under vacuum in the dark. The lipids were hydrated in 50 mM Tris/MES, pH 7.5, buffer to a final concentration of 20 mM, homogenized by freezing and thawing for 10 cycles, and stored in small aliquots at −80 °C. MSP1D1E3 was expressed and purified as previously described^39^ and dialyzed into 50 mM Tris/MES, pH 7.5, buffer. MSP1D1E3 was concentrated using a 10,000 MWCO filter concentrator and the final protein concentration was determined by A_280_ measurement (ε = 29,910 M^−1^⋅cm^−1^).

For reconstitution into nanodiscs, spin-labeled double-cysteine mutants in LMNG micelles were mixed with lipid mixture, MSP1D1E3, and sodium cholate in the following molar ratios: lipid:MSP1D1E3, 60:1; MSP1D1E3:*Ms*Spns, 10:1; and LMNG+cholate:lipid, 5:1. Reconstitution reactions were mixed at 4 °C for 1 h. Detergent was removed from the reaction by addition of 0.1 g/mL Biobeads (Bio-Rad) and incubation at 4 °C for 1 h. This was followed by another addition of 0.1 g/mL Biobeads with 1-h incubation, after which 0.2 mg/mL Biobeads were added and mixed overnight. The next day, 0.2 mg/mL Biobeads were added and mixed for 1 h. The reaction was filtered using a 0.45-µm filter to remove Biobeads. Full nanodiscs were separated from empty nanodiscs by size exclusion chromatography into 50 mM Tris/MES, pH 7.5, and 10% (vol/vol) glycerol buffer. The *Ms*Spns-containing nanodiscs were concentrated using Amicon ultra 100,000 MWCO filter concentrator, and then characterized using SDS/PAGE to verify reconstitution and estimate reconstitution efficiency. The concentration of spin-labeled mutants in nanodiscs was determined as described previously by comparing the intensity of the integrated continuous-wave electron paramagnetic resonance (CW-EPR) spectrum to that of the same mutant in detergent micelles^40^.

### Drug resistance assay

Resistance to toxic concentrations of capreomycin, conferred by *Ms*Spns WT and its mutants was carried out as previously described^7,28^. *Escherichia coli* BL21 (DE3) were transformed with empty pET19b vector, pET19b encoding *Ms*Spns WT, or mutant *Ms*Spns. A dense overnight culture from a single transformant was used to inoculate 10 mL of LB broth containing 0.1 mg/mL ampicillin to a starting Abs_600_ of 0.0375. Cultures were grown to Abs_600_ of 0.3 at 37 °C and expression of the encoded construct was induced with 1.0 μM IPTG (Gold Biotechnology). Expression was allowed to continue at 37 °C for 2 h, after which the Abs_600_ of the cultures was adjusted to 0.5. The cells were then used to inoculate (1:20 dilution, starting Abs_600_ = 0.025) a sterile 96-well microplate (Greiner) containing 50% LB broth, 0.1 mg/mL ampicillin, and 37.5 μg/mL of capreomycin. Microplates were incubated at 37 °C with shaking at 250 rpm for 2 to 8 h (6 h reported). The cell density (Abs_600_) was measured every 2 h on a SpectraMax i3 microplate reader and normalized to the 0 μg/mL drug well to obtain a relative absorbance, which accounts for growth behavior of the vector, WT, CL and other variants in the absence of drug. Experiments were performed at least in triplicate, and mean ± S.E.M. values were calculated; for *n* > 3, data were collected from multiple biological replicates. One-way ANOVA in OriginPro (OriginLab) showed that the population means of the *Ms*Spns mutants differed significantly at the 0.05 level, *F* (40, 234) = 28.81, *p* < 0.00001, whereas Tukey’s multiple-comparison test indicated that, in general, removal of the native cysteines and reintroduction of double-cysteine substitutions had little effect on the ability of *Ms*Spns to confer capreomycin resistance.

### Tryptophan fluorescence quenching

Purified *Ms*Spns in nanodisc buffer was adjusted to pH 4.0 or 9.0 using empirically determined volumes of citric acid or Tris, respectively, while maintaining equal protein concentrations across conditions. Samples were loaded into a quartz fluorescence cuvette (Starna Cells, catalog no. 16.40F-Q-10/Z15), and tryptophan fluorescence quenching was measured at 23 °C using a Horiba Scientific Fluoromax-4 spectrofluorometer. Excitation was at 295 nm, and emission was recorded from 310 to 370 nm. Fluorescence intensity at 335 nm was extracted from each spectrum to quantify the difference between pH 9.0 and pH 4.0 samples^28^. Experiments were performed in triplicate, and mean ± S.E.M. values were calculated; for *n* > 3, data were collected from multiple biological replicates. One-way ANOVA in OriginPro (OriginLab) showed that the population means of spin-labeled cysteine pairs were significantly different at the 0.05 level, *F* (40, 166) = 1160.9, *p* < 0.00001, whereas Tukey’s multiple-comparison test indicated that the constructs in nanodiscs did not differ significantly from CL *Ms*Spns.

### CW-EPR and DEER spectroscopy

CW-EPR spectra of spin-labeled *Ms*Spns samples were collected at room temperature on a Bruker EMX spectrometer operating at X-band frequency (9.5 GHz) using 10-mW incident power and a modulation amplitude of 1.6 G. DEER spectroscopy was performed on an Elexsys E580 EPR spectrometer operating at Q-band frequency (33.9 GHz) with the dead-time free four-pulse sequence at 83 K^41^. Pulse lengths were 20 ns (π/2) and 40 ns (π) for the probe pulses and 40 ns for the pump pulse. The frequency separation was 63 MHz. To ascertain the role of H^+^, samples were titrated to pH 4 and 9 with empirically determined amounts of 1 M citric acid and 1 M Tris, respectively, and confirmed by pH microelectrode (Mettler Toledo InLab Ultra-Micro-ISM) measurement. The substrate-bound state was generated by addition of 1 mM substrates at pH 7.5 or 9.0. Samples for DEER analysis were cryoprotected with 24% (vol/vol) glycerol and flash-frozen in liquid nitrogen.

Primary DEER decays were analyzed using home-written software operating in the Matlab (MathWorks) environment as previously described^42^. Briefly, the software carries out global analysis of the DEER decays obtained under different conditions for the same spin-labeled pair. The distance distribution is assumed to consist of a sum of Gaussians, the number and population of which are determined based on a statistical criterion. The generated confidence bands were determined from calculated uncertainties of the fit parameters. We also analyzed DEER decays individually and found that the resulting distributions agree with those obtained from global analysis. Comparison of the experimental distance distributions with the *Ms*Spns unbiased AF3 or DEER-guided refined models using a rotamer library approach was facilitated by the MMM software package^43^. Rotamer library calculations were conducted at 298 K.

### DEER-guided AlphaFold modeling

DEER-guided modeling of *Ms*Spns was performed using DEERFold^24^, a fine-tuned AlphaFold2-based^26^ network that incorporates experimental DEER distance distributions as pairwise distograms within the Evoformer module, together with the multiple sequence alignment (MSA), to bias structure prediction toward spin-label-constrained conformations. The DEERFold architecture, training strategy, and input-encoding scheme have been described previously^24^ and were used here without modification.

The *Ms*Spns amino acid sequence was used as input, and its MSA was generated with the ColabFold MSA pipeline^44^. For DEERFold predictions, the MSA was subsampled to an effective sequence depth of *N*_eff_ = 10^45^. For each restraint condition described below, 100 independent models were generated using distinct random seeds, producing a conformational ensemble for that condition. Models were ranked by the Earth Mover’s Distance (EMD) between the predicted and experimental distance distributions, and the highest-ranked subset, hereafter referred to as the “top models,” was retained for downstream structural analysis.

### Single-Gaussian restraints

For each spin pair, the experimental *P*(*r*) distribution measured under conditions favoring a single dominant conformation was approximated by a single Gaussian, defined by its mean distance and standard deviation. Restraints for the protonation-mimetic mutants E126Q and D38N were derived from the stabilized distance components observed under each condition. IF-state restraints were derived from distance distributions measured at pH 4.0, whereas OF-state restraints were derived from those measured at pH 9.0. Each single-Gaussian restraint set was supplied to DEERFold as an independent input, generating separate IF-, OF-, D38N-and E126Q-restrained ensembles. Because this restraint representation matches the input format on which DEERFold was fine-tuned^24^, it was expected to yield the closest agreement with the corresponding unbiased AlphaFold3 (AF3) reference models.

### Complete multi-Gaussian restraints

To determine whether the IF/OF preference was preserved with a less reduced representation of the experimental data, we also tested DEERFold using the full set of Gaussian components obtained from multi-Gaussian fitting of each DEER decay, thereby retaining the multi-peak character and breadth of the underlying *P*(*r*) distributions. Restraints were prepared independently for each DEER condition: pH 4.0, pH 7.5, and pH 9.0. Because DEERFold was fine-tuned exclusively on single-Gaussian distance distributions, these multi-Gaussian inputs fall outside the network’s training distribution and were therefore expected to generate more dispersed ensembles with reduced per-model agreement to the AF3 references. Nevertheless, this analysis provided an independent test of whether the underlying pH-dependent conformational preference was retained.

### Comparison with unbiased AF3 models

Unbiased reference models of *Ms*Spns in the IF and OF states were generated using AlphaFold3^27^, as implemented in the AlphaFold3 server. Five independent runs were performed for each state, and the highest-pLDDT model was selected as the reference conformation. DEERFold ensembles were then compared with the corresponding AF3 references using TM-score and Cα RMSD after structural superposition^46^. For each ensemble, we report the mean TM-score across the top models, together with the TM-score and Cα RMSD values for the single best-scoring model.

### DEER-guided refinement of the IF and OF conformations

The DEER distance constraints were used to refine the AF3-generated IF and OF models using a previously published approach^25^. Refinement was carried out iteratively in MODELLER^47^. In silico spin labeling was performed with the MMM software package^43^ using a rotamer-library approach. In each iteration, rotamer ensembles were first calculated at 298 K for the spin-labeled sites, and the rotamer that best matched the mean N–O midpoint position of the full ensemble was attached to the template structure provided to MODELLER. Refinement was performed using the centers of the Gaussian components corresponding to each conformational state, together with secondary-structure restraints derived from AlphaFold models. All refined models achieved a GA341 score of 1.0. After import into MMM, rotamer ensembles were recalculated (Supplementary Figs. 7 and 8), and models were ranked by the root-mean-square deviation between the experimental distance constraints and the corresponding model-derived distances across all restraints.

## Data availability

The generated data, including those from the DEER experiments, are available in the manuscript or the supplementary materials. The DEER data and generated models have been deposited to the Zenodo repository maintained by CERN, https://doi.org/10.5281/zenodo.20467517. Other data that support the findings of this study are available from the corresponding authors upon reasonable request.

## Acknowledgments

The authors wish to thank Dr. Ali Rasouli (Tajkhorshid Laboratory) for assistance with the molecular docking analysis. This work was supported by NIH grant R01-GM145783 and by a Lottie Hardy Foundation grant (to R.D.).

## Author contributions

R.D. designed the experiments. S.G., K.L.J., I.M., and K.D. expressed and purified the mutants and reconstituted in nanodiscs. R.D., and S.G. performed the EPR experiments and analyzed the data. T.W. and R.D. performed the computational modeling. K.L.J. performed the tryptophan fluorescence quenching experiments and cell growth assays. R.D., K.L.J., and S.G. wrote the paper. S.G. and K.L.J. contributed equally to this work; I.M. and T.W. also contributed equally.

## Supplementary Information

**This PDF file includes:**

Supplementary Figs. 1-15

---

**Supplementary Figure 1.**
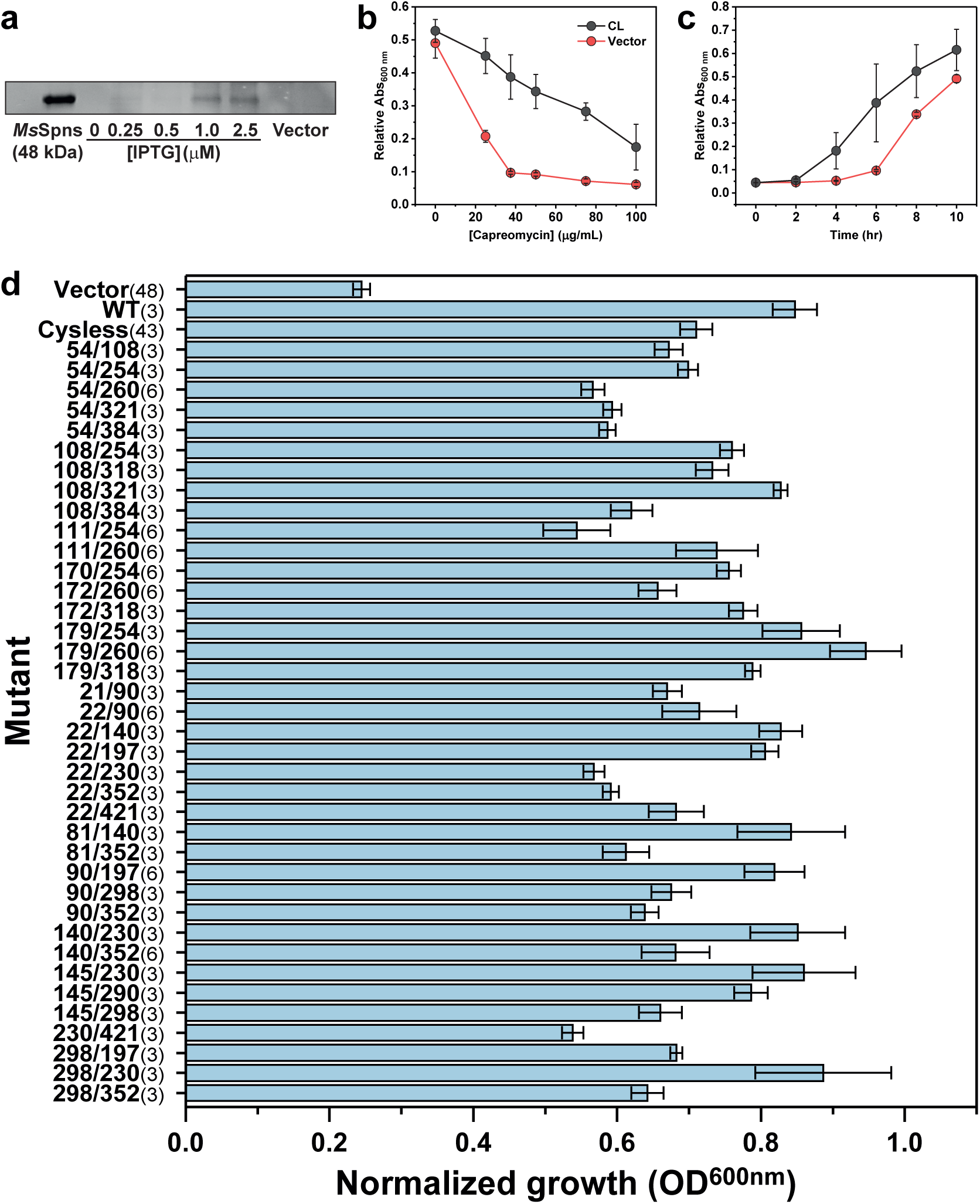
Capreomycin resistance profiles of *Ms*Spns mutants used for DEER spectroscopy. (**a**) Expression of cysteine-less (CL) *Ms*Spns in cell-growth assays at different IPTG concentrations (0–2.5 μM), visualized by SDS–PAGE and InVision His-tag staining. Purified CL *Ms*Spns is shown as a standard. 1 μM IPTG was used for protein expression in growth assays. (**b**) Expression of CL *Ms*Spns enhances cell survival at elevated capreomycin concentrations relative to the vector control. Measurements were performed in triplicate after 8 h at 37 °C. A_600nm_ values were normalized to the 0 μ g/mL capreomycin control, and standard deviations are shown. The optimal capreomycin concentration was 37.5 μg/mL. (**c**) Cells expressing CL *Ms*Spns show improved growth relative to the vector control at all time points in the presence of 37.5 μg/mL capreomycin. A_600nm_ values were normalized to the 0 μg/mL control; data points represent mean ± s.d. from triplicate measurements, as in b. (**d**) Growth of *Ms*Spns mutants at 37.5 μg/mL capreomycin, normalized to the 0 μg/mL control. Bars show mean ± S.E.M. for at least three replicates, with replicate numbers indicated in parentheses. For *n* > 3, data were collected from multiple biological replicates. One-way ANOVA showed significant differences among mutants at the 0.05 level *F*(40, 234) = 28.81, *p* < 0.00001, whereas Tukey’s multiple-comparison test indicated that the double-cysteine mutants did not differ significantly from WT or CL *Ms*Spns, showing that cysteine substitutions generally do not impair capreomycin resistance.

**Supplementary Figure 2.**
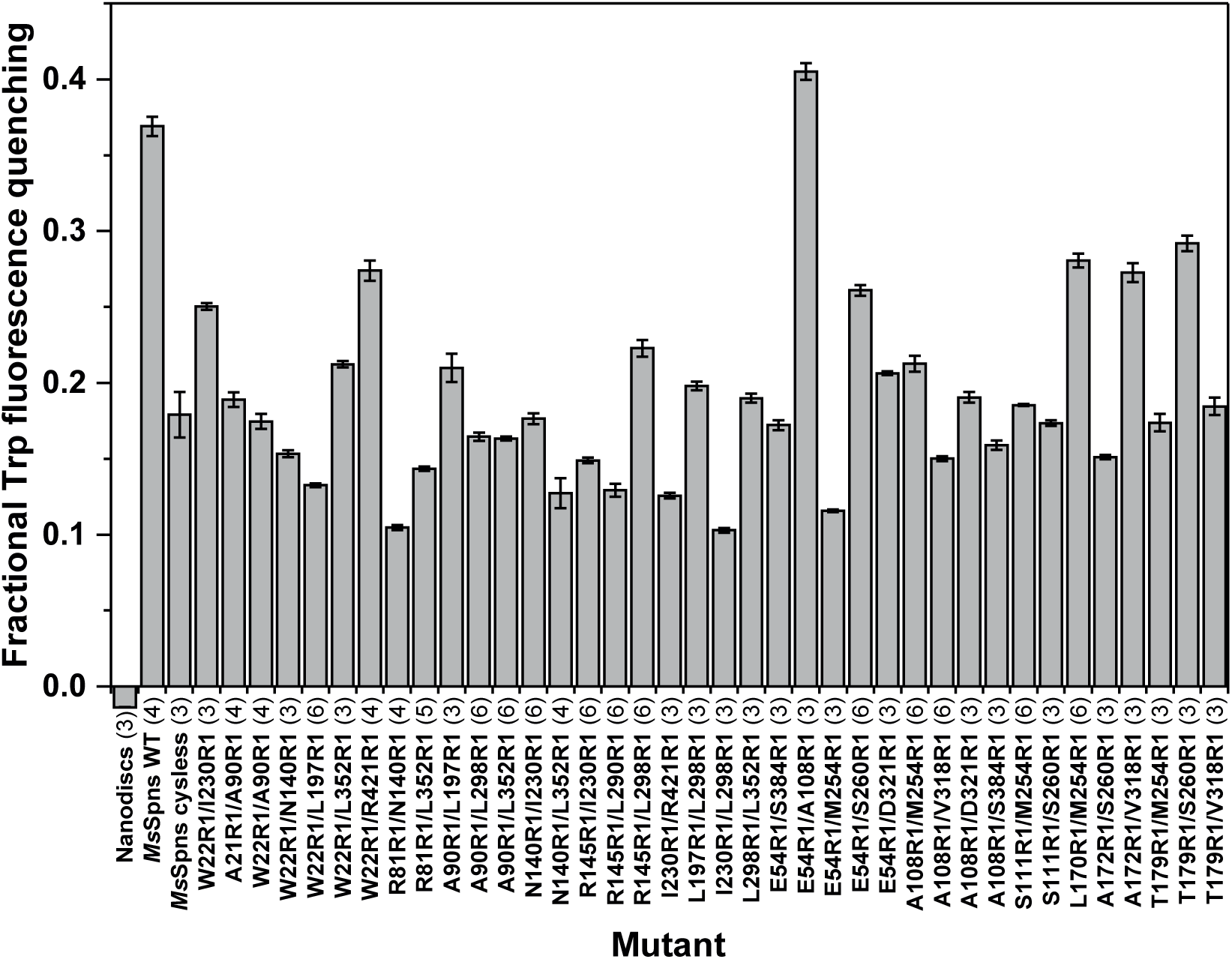
Fractional fluorescence quenching of *Ms*Spns mutants. Low-pH fluorescence quenching of native tryptophan residues was used as a surrogate reporter of conformational changes in spin-labeled *Ms*Spns reconstituted in nanodiscs, as described in Methods. Bars show mean ± S.E.M. for at least three replicates, with replicate numbers indicated in parentheses. For *n* > 3, data were collected from multiple biological replicates. One-way ANOVA showed significant differences among mutants *F*(40,166) = 1160.9, *p* < 0.00001, whereas Tukey’s multiple-comparison test indicated that removal of native cysteines, introduction of target-site cysteines, and spin labeling did not significantly alter conformationally coupled fluorescence quenching.

**Supplementary Figure 3.**
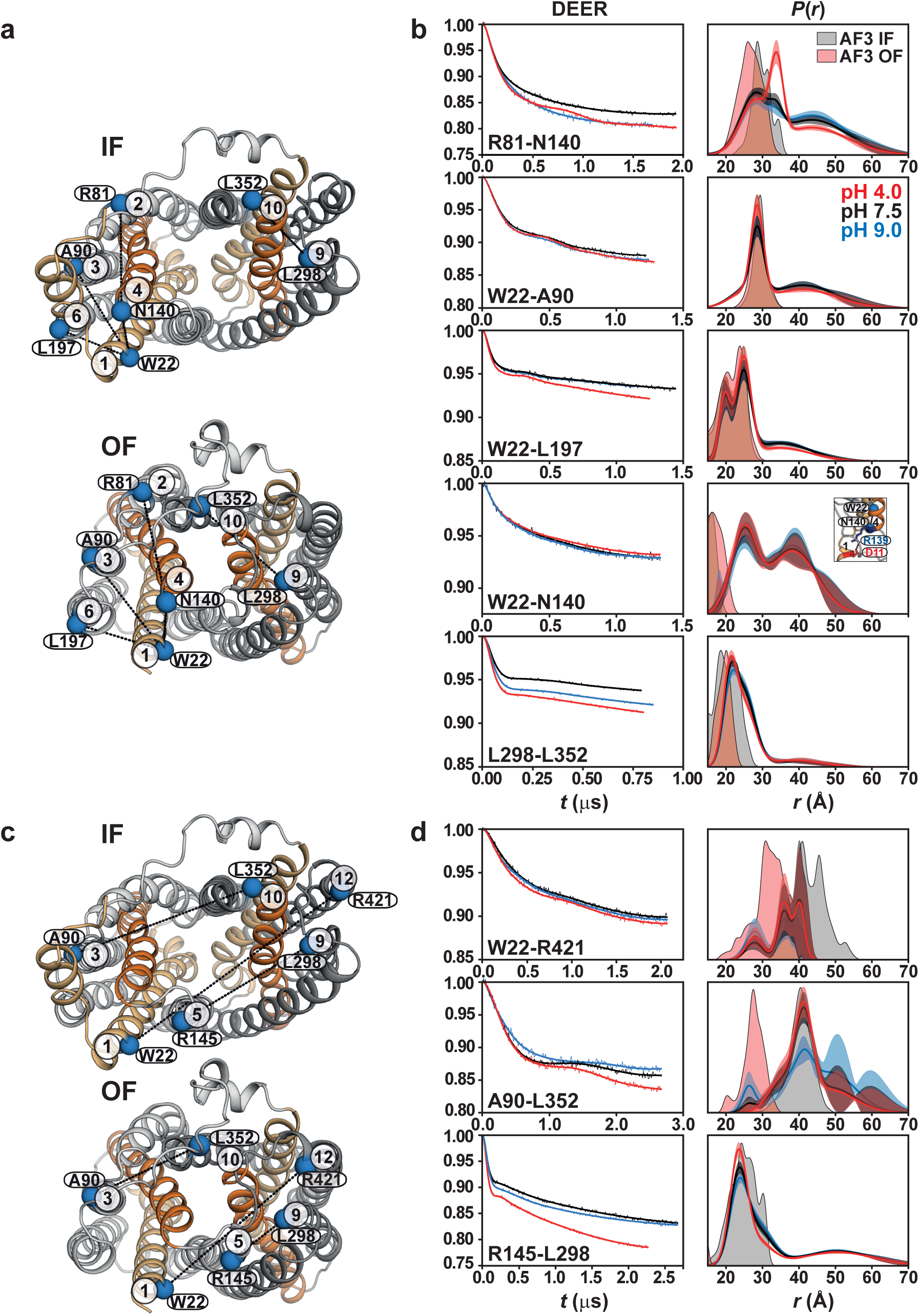
Intra-and interdomain DEER distance measurements on the intracellular side. (**a**) Intradomain spin-label pairs are shown as blue spheres on the intracellular side of the IF and OF models. (**b**) Raw DEER decays with fits (left) and corresponding distance distributions, *P*(*r*), measured in lipid nanodiscs (right). Distance distributions predicted from the IF and OF models are shown as gray and red shaded regions, respectively. Most intradomain pairs show limited proton-coupled conformational changes, except for R81–L352. (**c**) Interdomain spin-label pairs are shown as blue spheres on the intracellular side of the IF and OF models. (**d**) DEER decays with fits (left) and corresponding nanodiscs distance distributions, *P*(*r*) (right), with predicted IF and OF distributions shaded gray and red, respectively.

**Supplementary Figure 4.**
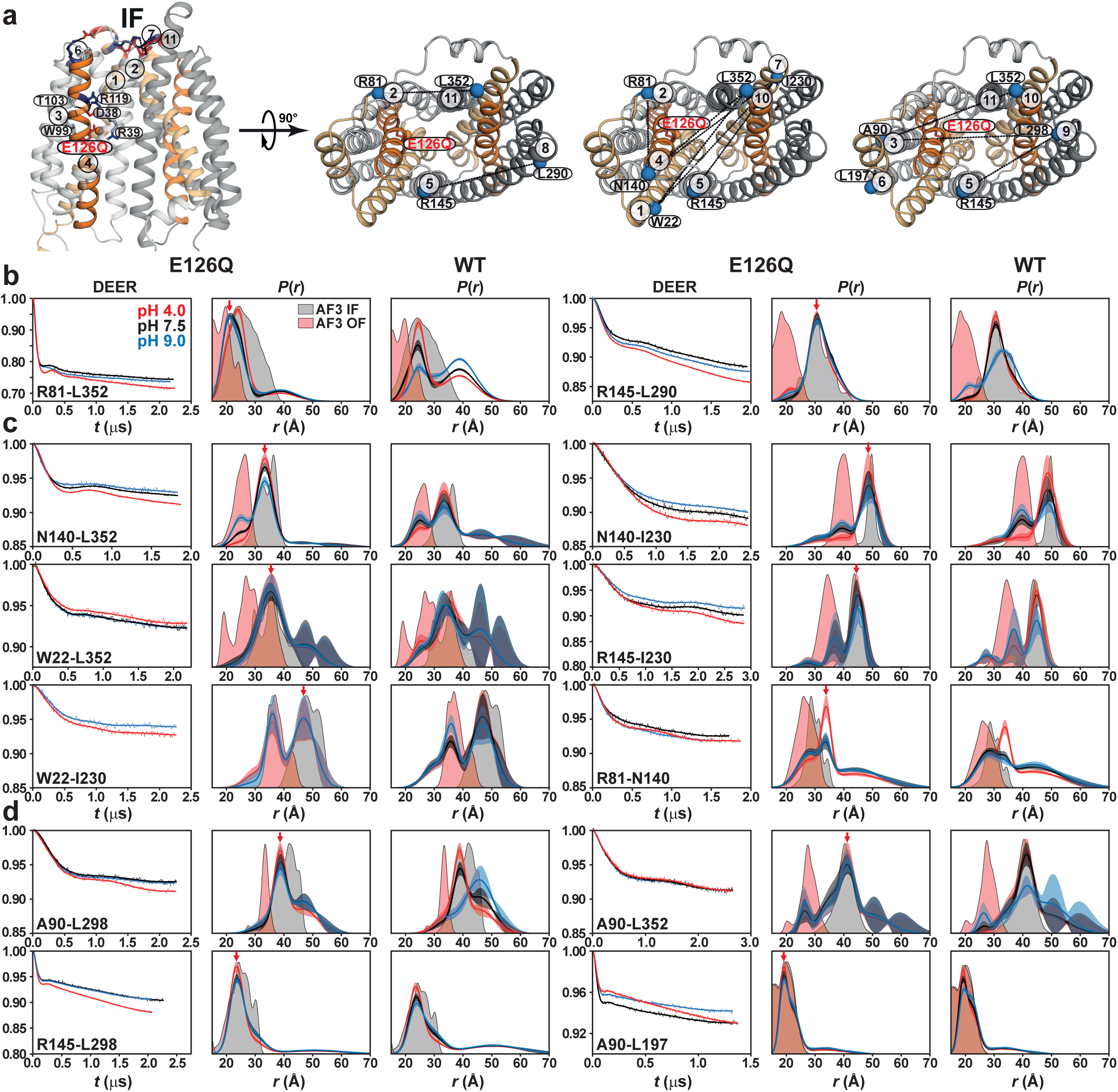
Glu126 protonation opens the intracellular side of *Ms*Spns with opposing effects on intracellular gates. (**a**) Spin-label pairs reporting on intracellular conformational changes are shown as blue spheres on the IF model. (**b**-**d**) Effects of the protonation-mimetic E126Q mutation on conformational equilibria at the intracellular gates (**b**), intracellular substrate-binding cavity (**c**) and support helices (**d**). The E126Q mutation was combined with the double-cysteine mutations. DEER measurements show that Glu126 protonation stabilizes an IF conformation, supporting its role as a protonation switch. Confidence bands represent 2σ uncertainty in *P*(*r*) associated with fitting of the primary DEER traces. Red arrows indicate the stabilized conformational state. Glu126 protonation closes the intracellular TM2/TM11 gate while opening the TM5/TM8 gate and expanding the substrate-binding cavity, thereby permitting substrate entry.

**Supplementary Figure 5.**
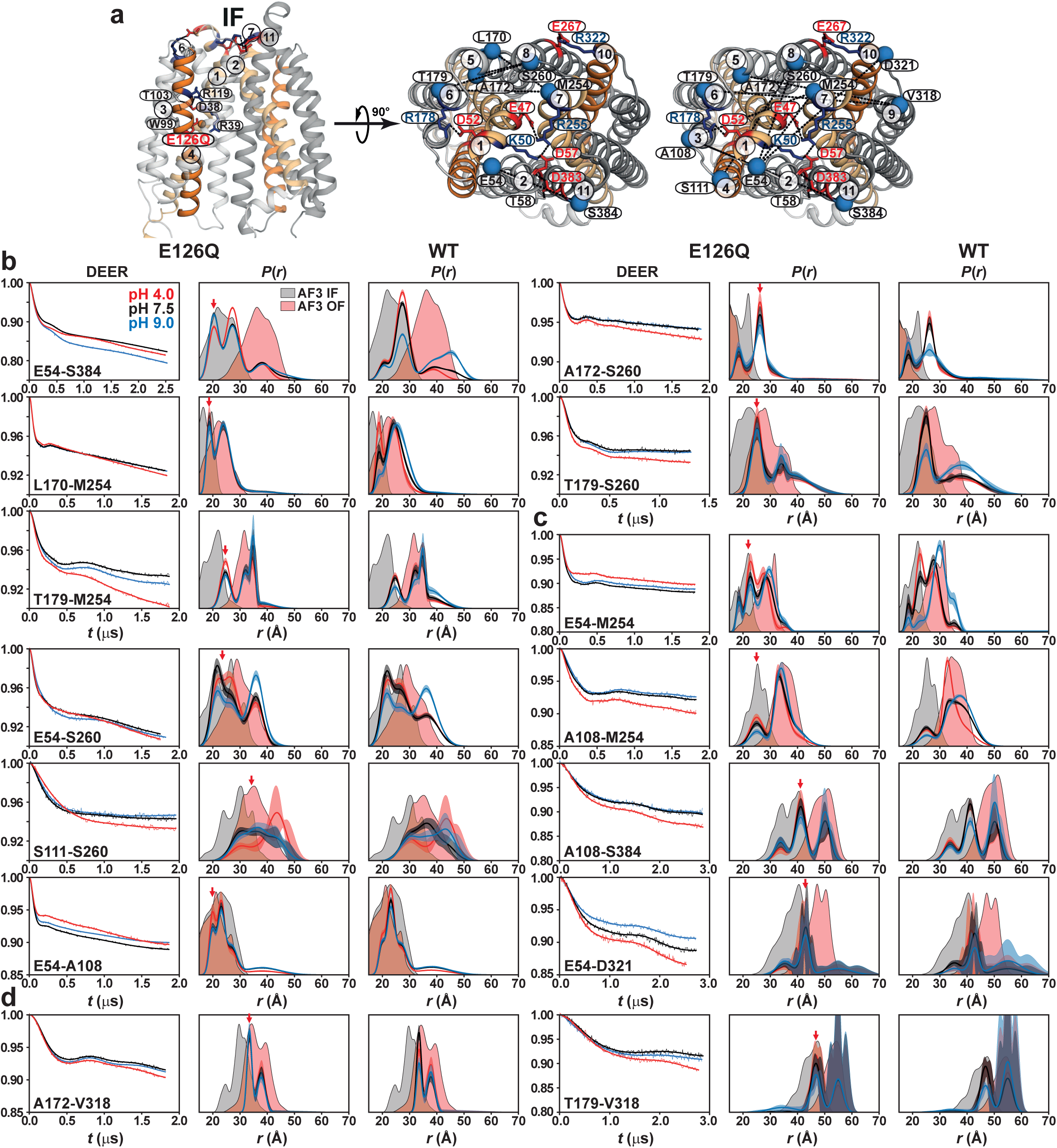
Glu126 protonation promotes extracellular closure in *Ms*5pns. (**a**) Spin-label pairs reporting on extracellular conformational changes are shown as blue spheres on the IF model. (**b**-**d**) Effects of the protonation-mimetic E126Q mutation on conformational equilibria at the extracellular gates (**b**), extracellular substrate-binding cavity (**c**) and support helices (**d**). The E126Q mutation was combined with the double-cysteine mutations. DEER measurements show that Glu126 protonation stabilizes an IF conformation, supporting its role as a protonation switch. Confidence bands represent 2σ uncertainty in *P*(*r*) associated with fitting of the primary DEER traces. Red arrows indicate the stabilized conformational state.

**Supplementary Figure 6.**
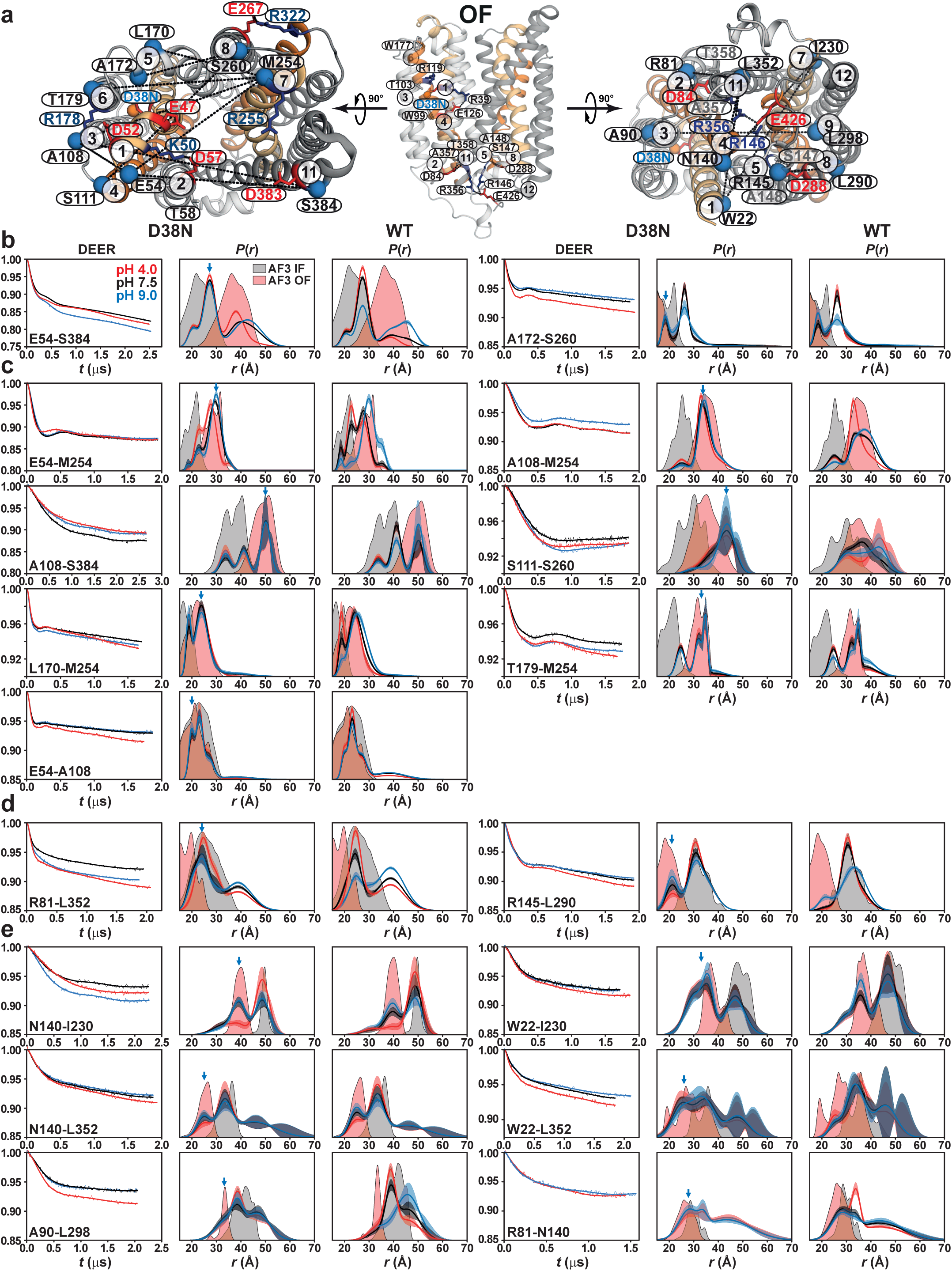
Asp38 protonation stabilizes the outward-facing conformation of *Ms*Spns with opposing effects on intracellular gates. (**a**) Spin-label pairs reporting on intracellular and extracellular conformational changes. (**b**-**e**) Effects of the protonation-mimetic D38N mutation on conformational equilibria at the extracellular (**b,c**) and intracellular (**d,e**) sides. DEER measurements show that Asp38 protonation stabilizes an OF conformation, supporting its role as a protonation switch. Blue arrows indicate the stabilized conformational state. In contrast to Glu126 protonation, Asp38 protonation closes the intracellular TM5/TM8 gate and substrate-binding cavity while opening the TM2/TM11 gate.

**Supplementary Figure 7.**
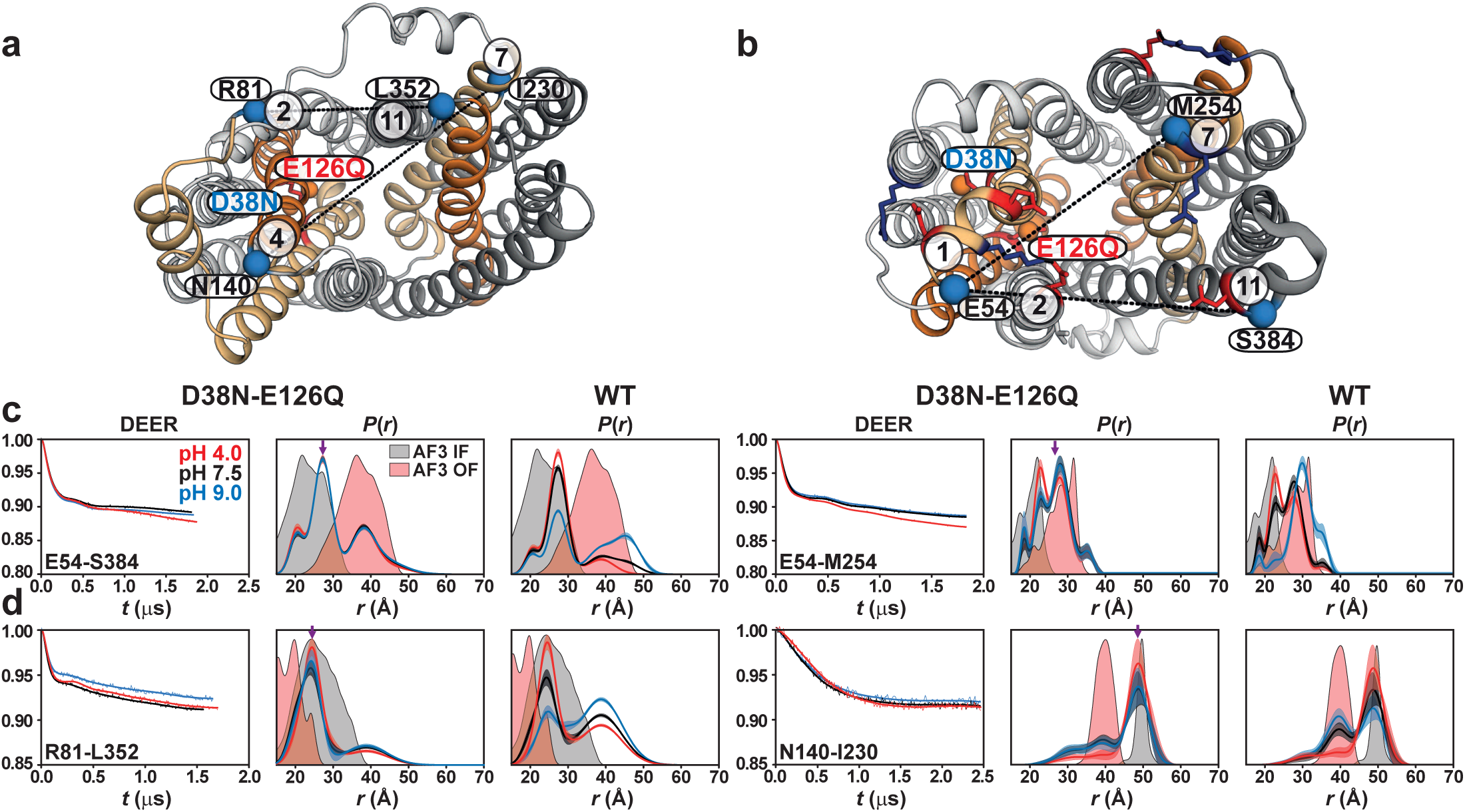
Asp38 and Glu126 protonation-mimetic substitutions shift *Ms*5pns toward an IF-like intermediate. (**a,b**) Spin-label pairs reporting on intracellular (**a**) and extracellular (**b**) conformational changes are shown as blue spheres on representative IF and OF *Ms*Spns models, respectively. (**c**) DEER analyses of extracellular reporters show that the D38N-E1260 double mutation shifts the TM2/TM11 gate pair E54-S384 and extracellular cavity pair E54-M254 toward shorter, more IF-like distance populations, particularly under acidic conditions, while retaining longer-distance OF-like states. (**d**) DEER analyses of intracellular reporters show a similar IF-like shift for the R81-L352 TM2/TM11 gate and N140-I230 cavity pairs. Together, these data indicate that combined Asp38 and Glu126 protonation does not lock *Ms*Spns into a single canonical IF or OF state, but instead biases the ensemble toward an IF-like intermediate. These results are consistent with opposing, region-specific contributions of Asp38 and Glu126 to intracellular and extracellular conformational equilibria. Purple arrows mark mutation-enriched distance populations.

**Supplementary Figure 8.**
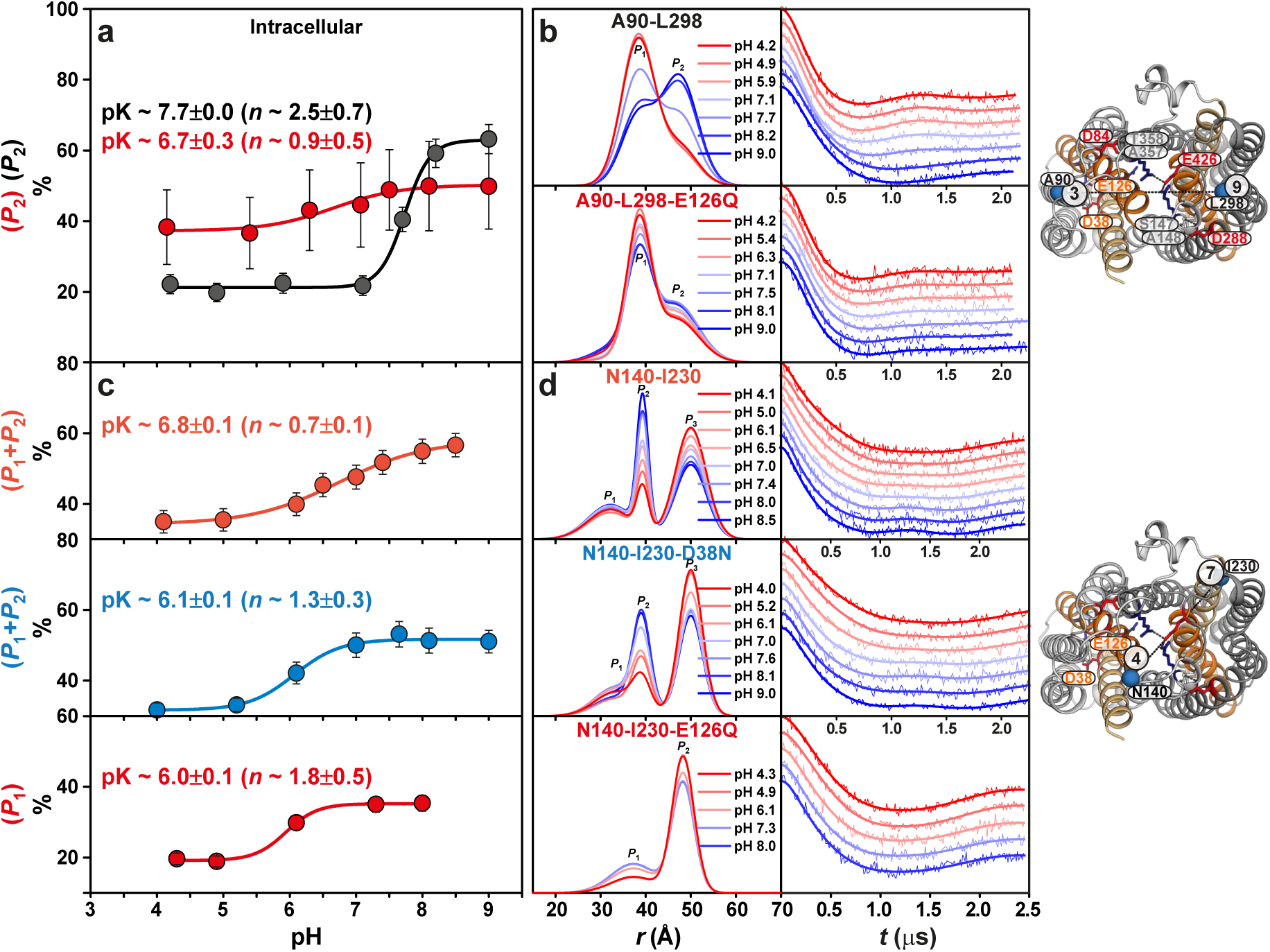
Glu126 protonation regulates intracellular pH-dependent conformational changes, whereas Asp38 modestly affects the substrate-binding cavity. (**a,b**) Baseline-corrected and normalized DEER traces with fits, corresponding distance distributions, and pH-dependent population changes for the intracellular support-helix pair A90–L298, with and without the protonation-mimetic E126Q mutation. Population changes in the increasing-distance peaks were used to estimate apparent pK values for conformational transitions. The Glu126 protonation-mimetic mutation markedly attenuates pH-dependent rearrangement of the support helices. (**c,d**) Corresponding analysis of the intracellular substrate-binding cavity reporter N140–I230, with and without D38N or E126Q. The substrate-binding cavity is only modestly altered by D38N but is more strongly affected by E126Q, indicating a dominant role for Glu126 in regulating the intracellular cavity. Error bars in **a** and **c** represent 2σ (95%) confidence intervals for the fitted peak populations shown in **b** and **d**, respectively.

**Supplementary Figure 9.**
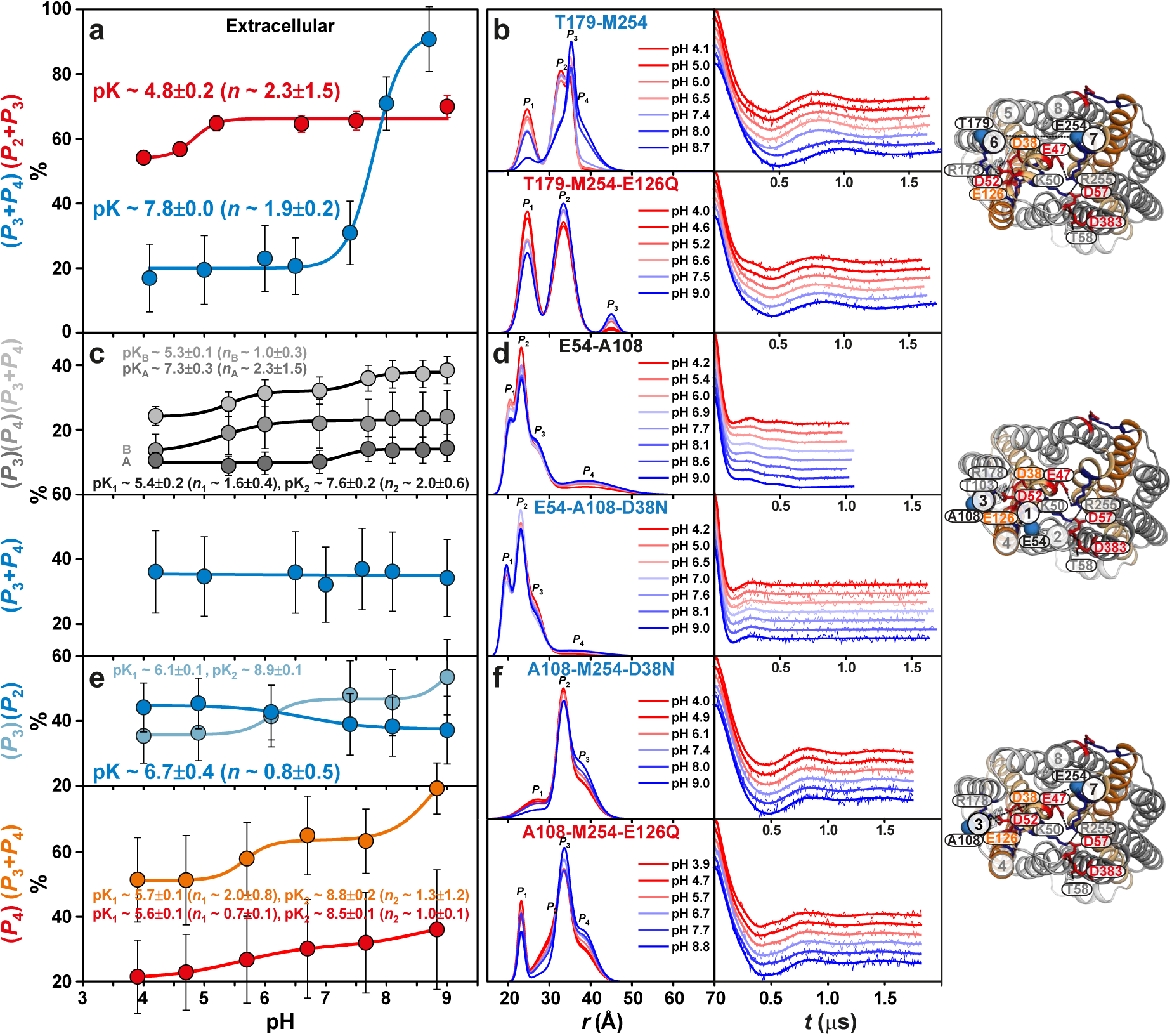
Asp38 and Glu126 regulate extracellular pH-dependent conformational changes in *Ms*Spns. (**a,b**) Baseline-corrected and normalized DEER traces with fits, corresponding distance distributions, and pH-dependent population changes for the extracellular T179–M254 pair, which indirectly reports on the TM5/8 lateral gate, with and without the protonation-mimetic E126Q mutation. Population changes in the increasing-distance peaks were used to estimate apparent pK values for extracellular conformational transitions. As on the intracellular side, the extracellular gate and support helices are strongly modulated by Glu126 protonation. (**c,d**) Analysis of the extracellular TM1/3 pair E54–A108, which probes opening of the putative periplasmic proton-translocation pathway along TMs 1, 3 and 6, with and without D38N. D38N strongly suppresses pH-dependent conformational changes, supporting a dominant role for Asp38 in regulating this pathway. (**e,f**) Extracellular TM3/7 pair A108–M254, which reports on the extracellular substrate-binding cavity, with D38N or E126Q. D38N largely suppresses, whereas E126Q attenuates, pH-dependent conformational changes in this extracellular reporter (wildtype: Fig. 3d). Error bars in **a, c** and **e** represent 2σ (95%) confidence intervals for the fitted peak populations shown in **b, d** and **f**, respectively.

**Supplementary Figure 10.**
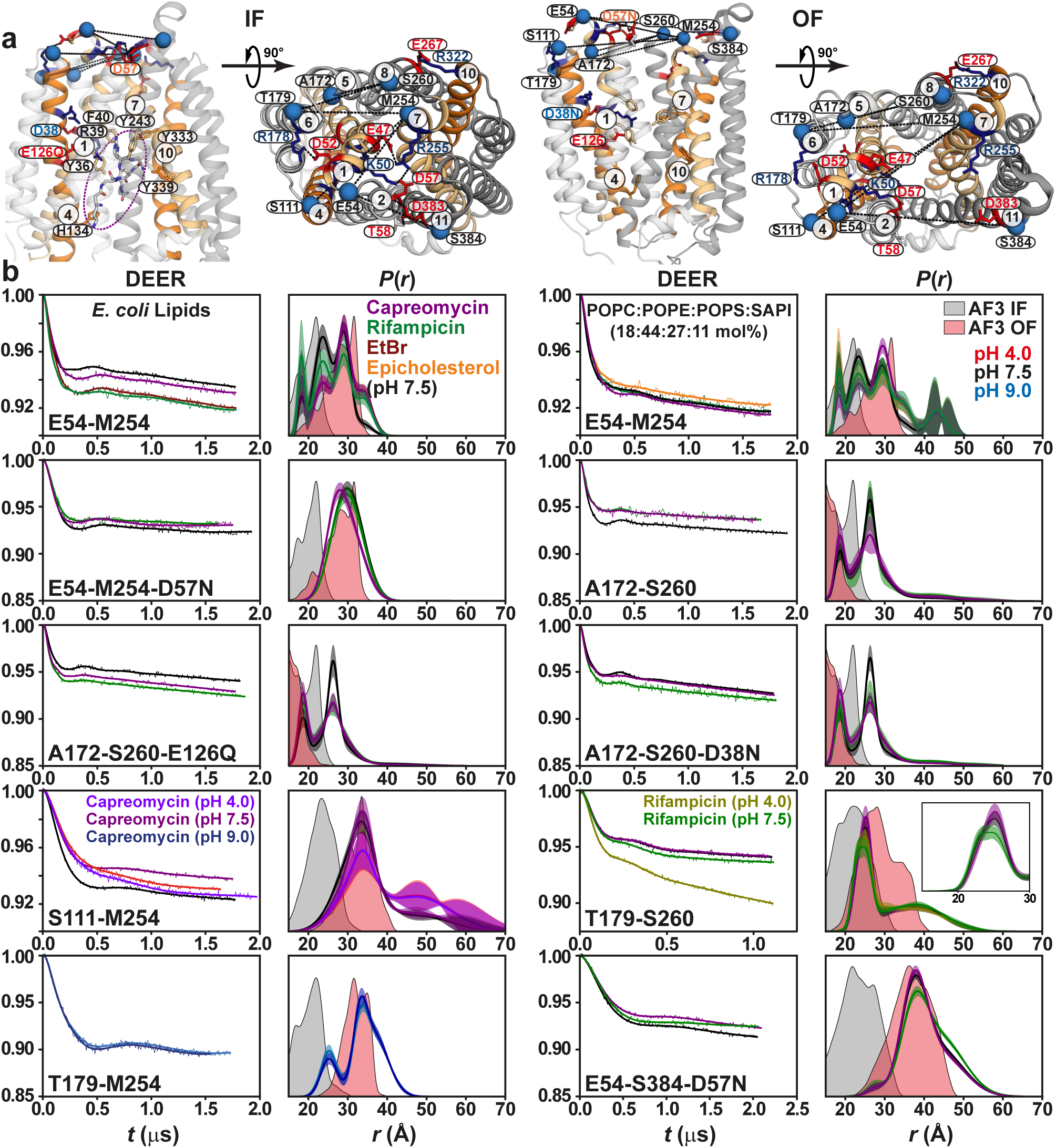
Hydrophobic and hydrophilic substrates differentially shift extracellular *Ms*Spns conformational equilibria. (**a**) Membrane and periplasmic views of the IF and OF models. Capreomycin (circled) is docked in the IF substrate-binding cavity, with conserved aromatic residues in binding cavity shown as sticks and DEER spin-label pairs shown as blue spheres. Functional residue interactions stabilizing the IF conformation are highlighted in red and blue in the periplasmic views. (**b**) Raw DEER decays with fits and corresponding extracellular distance distributions measured in the apo state and in the presence of substrates, with and without protonation-mimetic mutations. Two lipid compositions were tested for the E54-M254 pair. Water-soluble cationic substrates, including capreomycin and ethidium bromide, stabilize the OF conformation, whereas lipophilic substrates, including rifampicin and epicholesterol, shift the equilibrium toward the IF state.

**Supplementary Figure 11.**
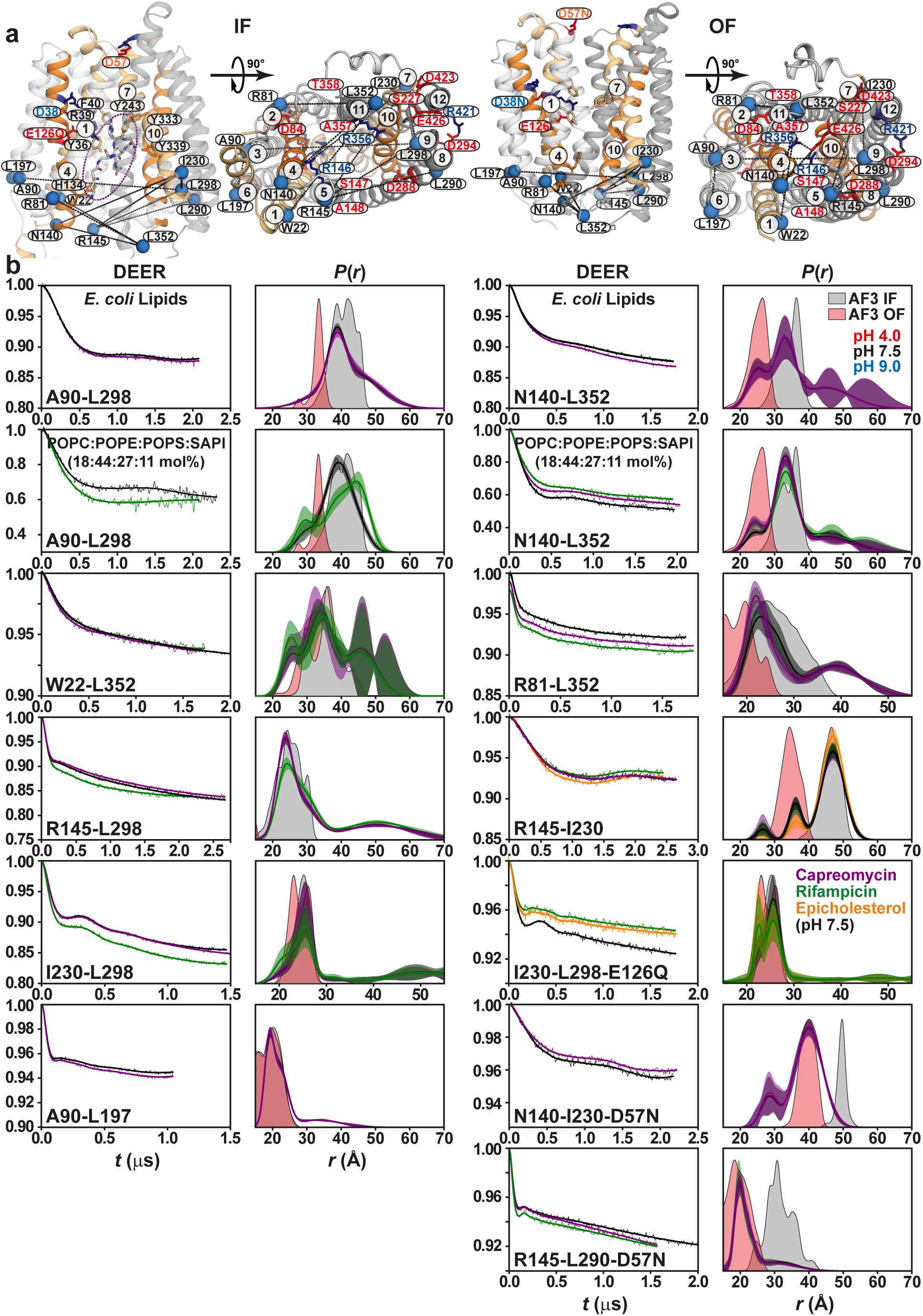
Substrate hydrophobicity drives opposing shifts in intracellular *Ms*Spns conformational equilibria. (**a**) Membrane and cytoplasmic views of the IF and OF models. Capreomycin is docked in the IF substrate-binding cavity, with conserved aromatic residues shown as sticks and DEER spin-label pairs shown as blue spheres. Functional residue interactions that stabilize the OF conformation are highlighted in red and blue in the cytoplasmic views. (**b**) Raw DEER decays with fits and corresponding intracellular distance distributions measured in the apo state and in the presence of substrates, with and without protonation-mimetic mutations. Two lipid compositions were tested for the A90-L298 and N140-L352 pairs. Water-soluble cationic substrates, including capreomycin and ethidium bromide, s tabilize the OF conformation, whereas lipophilic substrates, including rifampicin and epicholesterol, shift the equilibrium toward the IF state.

**Supplementary Figure 12.**
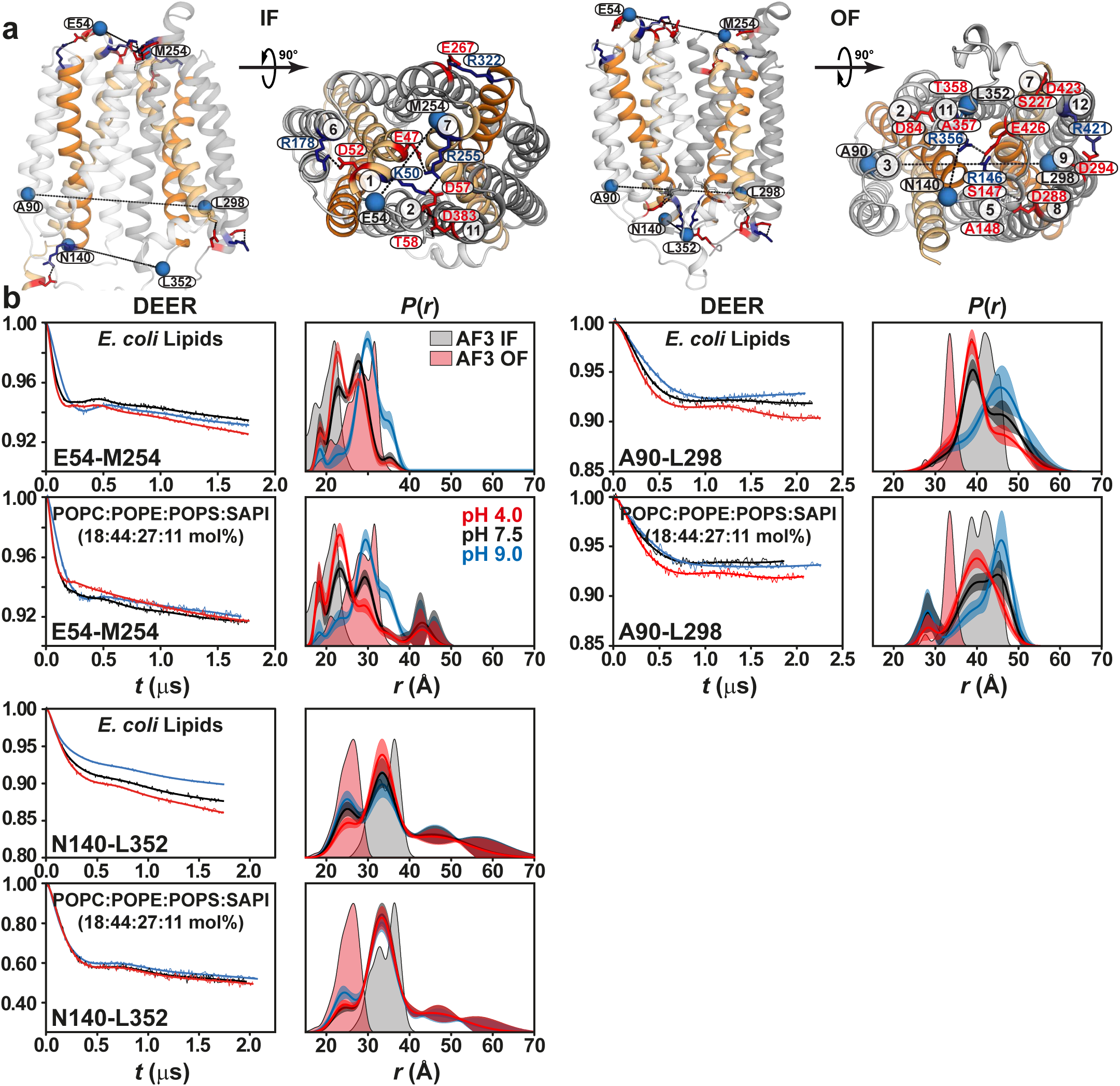
Lipid composition modulates *Ms*Spns conformational equilibria. (**a**) Membrane and periplasmic views of the IF model and membrane and cytoplasmic views of the OF model. DEER spin-label pairs are shown as blue spheres. Functional residue interactions, including the intracellular and extracellular proton-sensing networks, are shown as red and blue sticks and labels. (**b**) Raw DEER decays with fits and corresponding distance dis1tributions measured in the apo state for two lipid compositions, using the extracellular pair E54-M254 and the intracellular pairs A90-L298 and N140-L352. The lipid composition containing L-α-phosphatidylinositol (SAPI) shifts the equilibrium toward the IF state across protonation conditions, resembling the effect of lipophilic subs trates such as rifampicin and sugges ting possible phospholipid subs trate recognition or allos teric s tabilization

**Supplementary Figure 13.**
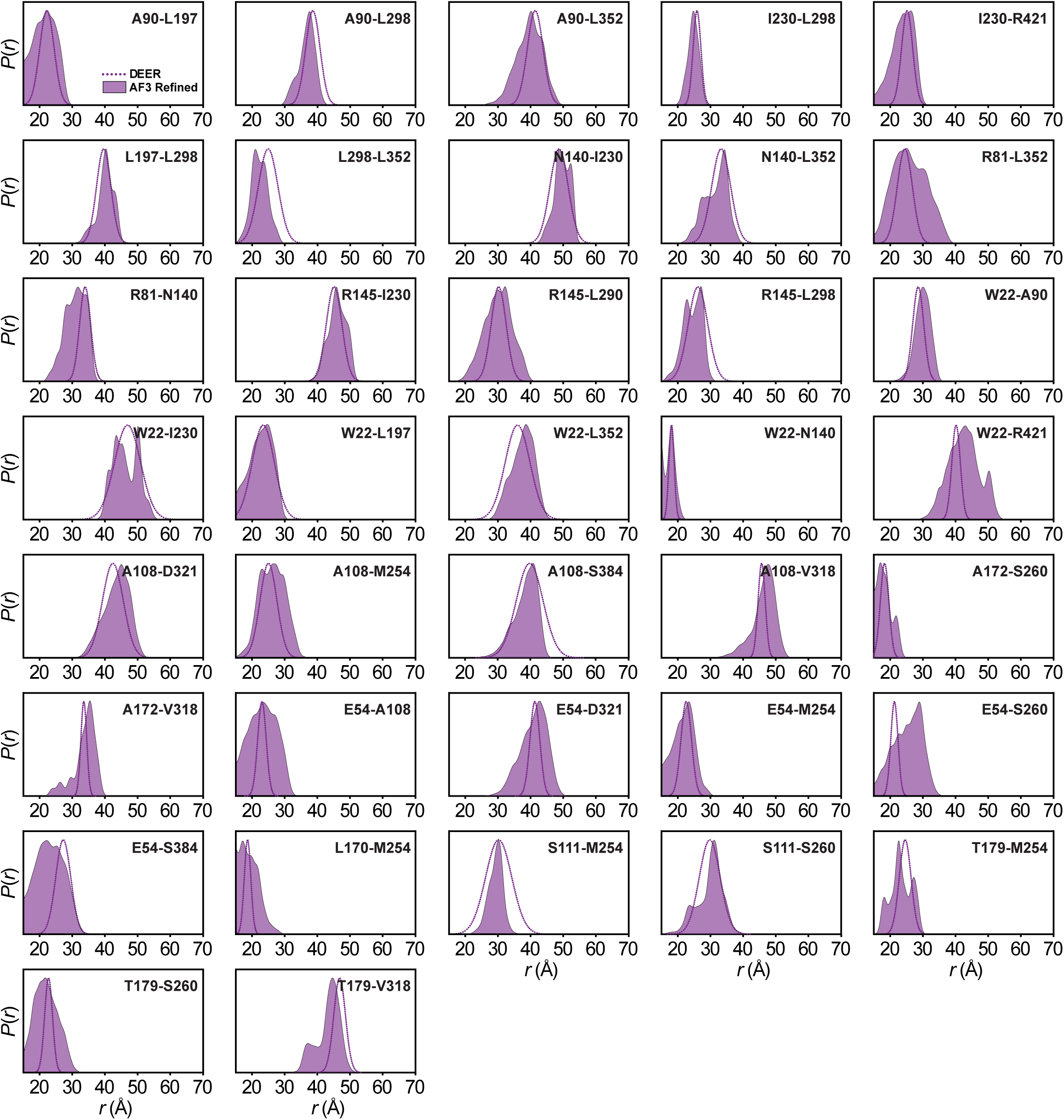
DEER-guided refinement of the IF *Ms*Spns AlphaFold3 model. Distance distributions predicted from the refined IF model (purple shading) are compared with IF-associated experimental DEER distance distributions measured in nanodiscs (dotted purple lines). Overall, the refined model closely captures the experimental distance subpopulations.

**Supplementary Figure 14.**
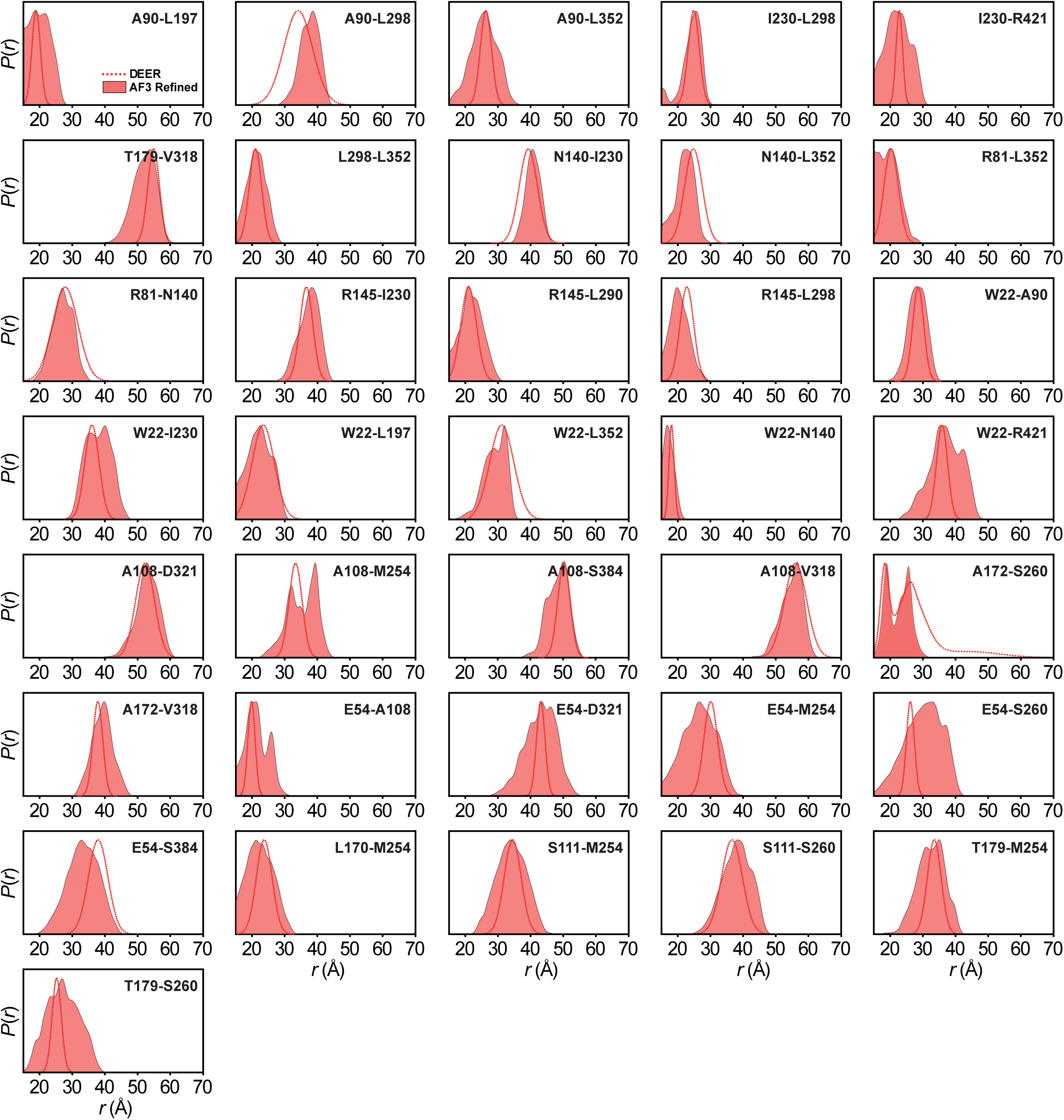
DEER-guided refinement of the OF *Ms*Spns AlphaFold3 model. Dis1tance distributions predicted from the refined OF model (red shading) are compared with OF-associated experimental DEER distance distributions measured in nanodiscs (dotted red lines). Overall, the refined model closely captures the experimental distance subpopulations.

**Supplementary Figure 15.**
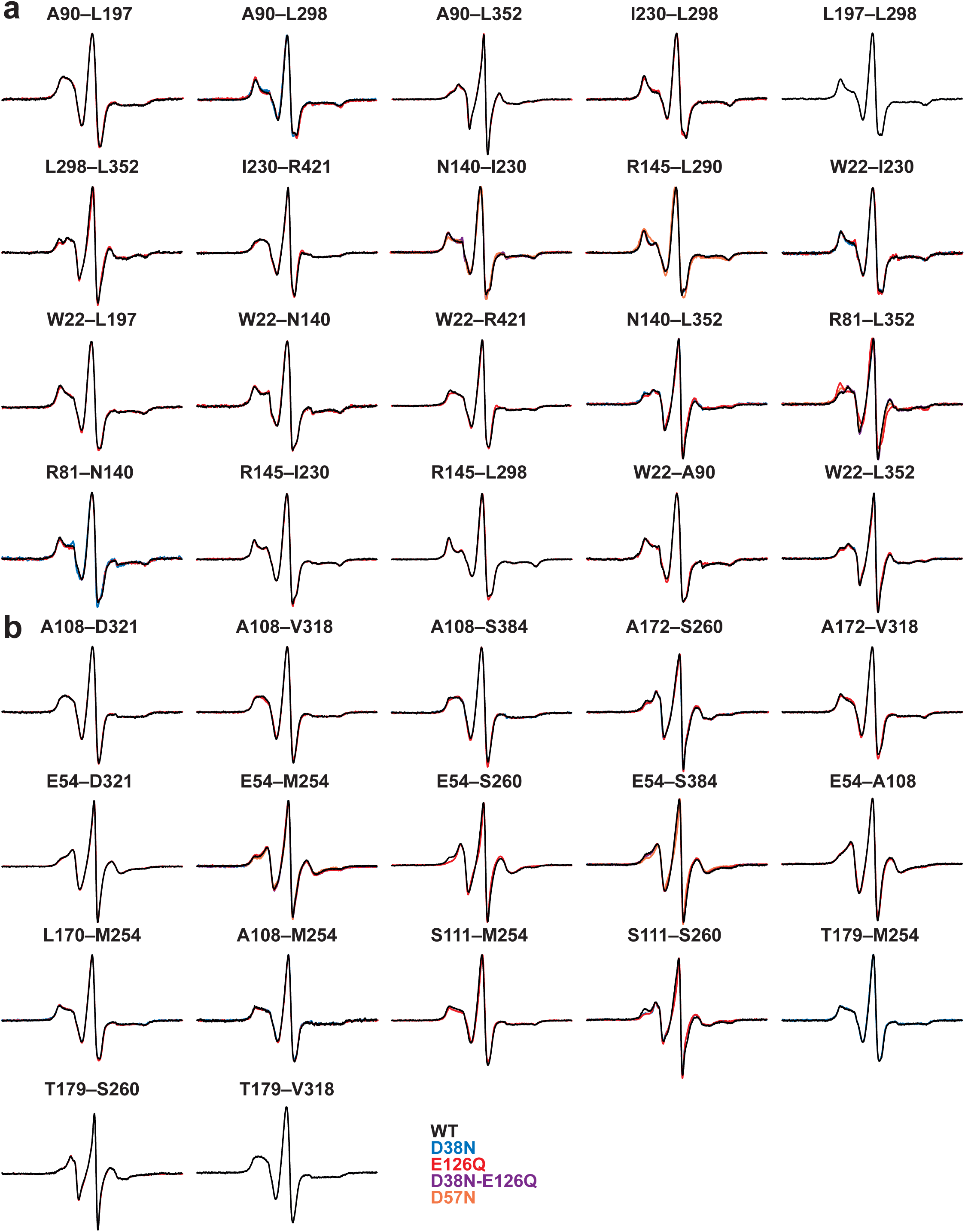
CW EPR spectra of spin-labeled double-cysteine mutants for DEER spectroscopy.

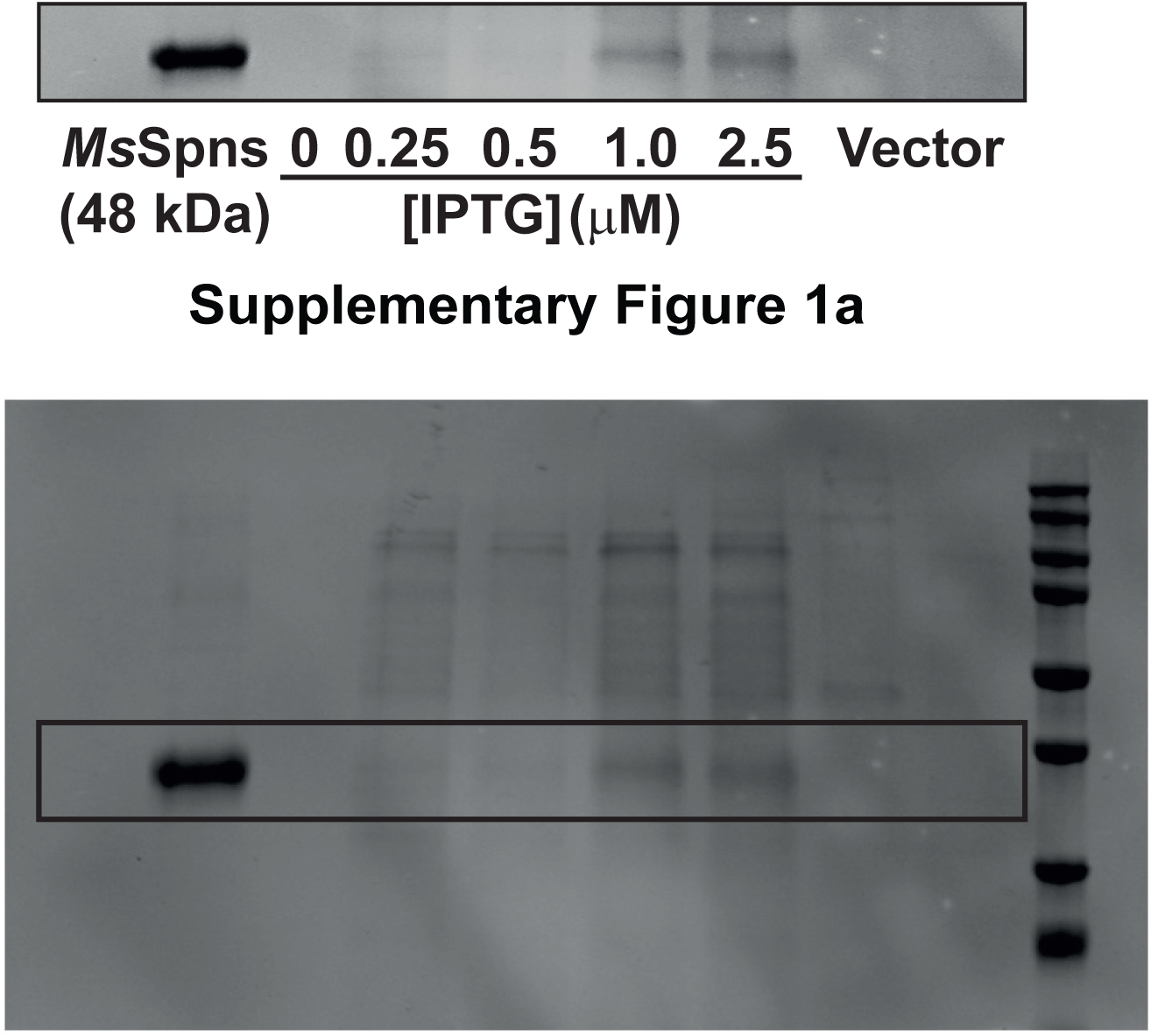

## References

1. van der Weyden, L. et al. Genome-wide in vivo screen identifies novel host regulators of metastatic colonization. Nature 541, 233–236 (2017).

2. Chen, H. et al. Structural and functional insights into Spns2-mediated transport of sphingosine-1-phosphate. Cell 186, 2644–2655 (2023).

3. Pang, B. et al. Molecular basis of Spns2-facilitated sphingosine-1-phosphate transport. Cell Research 34, 173–176 (2024).

4. Duan, Y. N. et al. Structural basis of Sphingosine-1-phosphate transport via human SPNS2. Cell Research 34, 177–180 (2024).

5. Tang, H.P. et al. The solute carrier SPNS2 recruits PI(4,5)P_2_ to synergistically regulate transport of sphingosine-1-phosphate. Molecular Cell 83, 2739–2752 (2023).

6. Li, H.Z. et al. Transport and inhibition of the sphingosine-1-phosphate exporter SPNS2. Nature Communications 16, 721 (2025).

7. Dastvan, R. et al. Proton-driven alternating access in a spinster lipid transporter. Nature Communications 13, 5161 (2022).

8. He, M.L. et al. Spns1 is a lysophospholipid transporter mediating lysosomal phospholipid salvage. Proceedings of the National Academy of Sciences of the USA 119, e2210353119 (2022).

9. Ha, H.T.T., et al. Lack of SPNS1 results in accumulation of lysolipids and lysosomal storage disease in mouse models. Jci Insight 9, e175462 (2024).

10. Chen, H. et al. Molecular basis of Spns1-mediated lysophospholipid transport from the lysosome. Proceedings of the National Academy of Sciences of the USA 122, e2409596121 (2025).

11. Spiegel, S. et al. New insights into functions of the sphingosine-1-phosphate transporter SPNS2. Journal of Lipid Research 60, 484–489 (2019).

12. He, M.L. et al. SPNS1 variants cause multiorgan disease and implicate lysophospholipid transport as critical for mTOR-regulated lipid homeostasis. The Journal of Clinical Investigation 135, e193099 (2025).

13. Zhou, F. et al. Crystal structure of a bacterial homolog to human lysosomal transporter, spinster. Science Bulletin 64, 1310–1317 (2019).

14. Weigel, C. et al. SPNS2 exports sphingosine-1-phosphate and imports glucose. Nature Communications 17, 3901 (2026).

15. Zhang, Z., Wang, R. & Xie, J.P. *Mycobacterium smegmatis* MSMEG_3705 encodes a selective major facilitator superfamily efflux pump with multiple roles. Current Microbiology 70, 801–809 (2015).

16. Zhang, X.J.C. et al. Energy coupling mechanisms of MFS transporters. Protein Science 24, 1560–1579 (2015).

17. Drew, D. et al. Structures and General Transport Mechanisms by the Major Facilitator Superfamily (MFS). Chemical Reviews 121, 5289–5335 (2021).

18. Cater, R.J. et al. Structural basis of omega-3 fatty acid transport across the blood-brain barrier. Nature 595, 315–319 (2021).

19. Radestock, S. & Forrest, L.R. The Alternating-Access Mechanism of MFS Transporters Arises from Inverted-Topology Repeats. Journal of Molecular Biology 407, 698–715 (2011).

20. Madej, M.G. et al. Evolutionary mix-and-match with MFS transporters. Proceedings of the National Academy of Sciences of the USA 110, 5870–5874 (2013).

21. Schiemann, O. et al. Benchmark Test and Guidelines for DEER/PELDOR Experiments on Nitroxide-Labeled Biomolecules. Journal of the American Chemical Society 143, 17875–17890 (2021).

22. Verhalen, B. et al. Energy transduction and alternating access of the mammalian ABC transporter P-glycoprotein. Nature 543, 738–741 (2017).

23. Dastvan, R. & Stoll, S. Recent advances in quantifying protein conformational ensembles with dipolar EPR spectroscopy. Current Opinion in Structural Biology 94, 103139 (2025).

24. Wu, T. et al. Modeling protein conformational ensembles by guiding AlphaFold2 with Double Electron Electron Resonance (DEER) distance distributions. Nature Communications 16, 7107 (2025).

25. Dastvan, R. et al. Protonation-dependent conformational dynamics of the multidrug transporter EmrE. Proceedings of the National Academy of Sciences of the USA 113, 1220–1225 (2016).

26. Jumper, J. et al. Highly accurate protein structure prediction with AlphaFold. Nature 596, 583–589 (2021).

27. Abramson, J. et al. Accurate structure prediction of biomolecular interactions with AlphaFold 3. Nature 630, 493–500 (2024).

28. Jagessar, K.L., Mchaourab, H.S. & Claxton, D.P. The N-terminal domain of an archaeal multidrug and toxin extrusion (MATE) transporter mediates proton coupling required for prokaryotic drug resistance. Journal of Biological Chemistry 294, 12807–12814 (2019).

29. Jagessar, K.L. et al. Sequence and structural determinants of ligand-dependent alternating access of a MATE transporter. Proceedings of the National Academy of Sciences of the USA 117, 4732–4740 (2020).

30. Jiang, D.H. et al. Structure of the YajR transporter suggests a transport mechanism based on the conserved motif A. Proceedings of the National Academy of Sciences of the USA 110, 14664–14669 (2013).

31. Nandakumar, M., Nathan, C. & Rhee, K.Y. Isocitrate lyase mediates broad antibiotic tolerance in *Mycobacterium tuberculosis*. Nature Communications 5, 4306 (2014).

32. Jackson, M., Crick, D.C. & Brennan, P.J. Phosphatidylinositol is an essential phospholipid of mycobacteria. Journal of Biological Chemistry 275, 30092–30099 (2000).

33. Dufrisne, M.B. et al. Structural and Functional Characterization of Phosphatidylinositol-Phosphate Biosynthesis in Mycobacteria. Journal of Molecular Biology 432, 5137–5151 (2020).

34. Martens, C. et al. Direct protein-lipid interactions shape the conformational landscape of secondary transporters. Nature Communications 9, 4151 (2018).

35. Szafraniec, M. et al. Capreomycin and hygromycin B modulate the catalytic activity of the delta ribozyme in a manner that depends on the protonation and complexation with Cu ions of these antibiotics. Dalton Transactions 41, 9728–9736 (2012).

36. Wisedchaisri, G. et al. Proton-coupled sugar transport in the prototypical major facilitator superfamily protein XylE. Nature Communications 5, 4521 (2014).

37. Leano, J.B. et al. Structures suggest a mechanism for energy coupling by a family of organic anion transporters. Plos Biology 17, e3000260 (2019).

38. Dmitrieva, N. et al. Transport mechanism of DgoT, a bacterial homolog of SLC17 organic anion transporters. EMBO Journal 241, 6740–6765 (2024).

39. Mishra, S. et al. Conformational dynamics of the nucleotide binding domains and the power stroke of a heterodimeric ABC transporter. Elife 3, e02740 (2014).

40. Zou, P. & Mchaourab, H.S. Increased Sensitivity and Extended Range of Distance Measurements in Spin-Labeled Membrane Proteins: Q-Band Double Electron-Electron Resonance and Nanoscale Bilayers. Biophysical Journal 98, L18–L20 (2010).

41. Jeschke, G. DEER distance measurements on proteins. Annual Review of Physical Chemistry 63, 419–446 (2012).

42. Hustedt, E.J., Stein, R.A. & Mchaourab, H.S. Protein functional dynamics from the rigorous global analysis of DEER data: Conditions, components, and conformations. The Journal of General Physiology 153, e201711954 (2021).

43. Polyhach, Y., Bordignon, E. & Jeschke, G. Rotamer libraries of spin labelled cysteines for protein studies. Physical Chemistry Chemical Physics 13, 2356–2366 (2011).

44. Mirdita, M. et al. ColabFold: making protein folding accessible to all. Nature Methods 19, 679–682 (2022).

45. Wu, T.Q. et al. Analysis of several key factors influencing deep learning-based inter-residue contact prediction. Bioinformatics 36, 1091–1098 (2020).

46. Zhang, Y. & Skolnick, J. Scoring function for automated assessment of protein structure template quality. Proteins 57, 702–710 (2004).

47. Sali, A. & Blundell, T.L. Comparative Protein Modeling by Satisfaction of Spatial Restraints. Journal of Molecular Biology 234, 779–815 (1993).

48. Berka, K. et al. MOLEonline 2.0: interactive web-based analysis of biomacromolecular channels. Nucleic Acids Research 40, W222–W227 (2012).

